# Rapid UPF1 depletion illuminates the temporal dynamics of the NMD-regulated transcriptome in human cells

**DOI:** 10.1101/2024.03.04.583328

**Authors:** Volker Boehm, Damaris Wallmeroth, Paul O. Wulf, Luiz Gustavo Teixeira Alves, Oliver Popp, Maximilian Riedel, Emanuel Wyler, Marek Franitza, Jennifer V. Gerbracht, Kerstin Becker, Karina Polkovnychenko, Simone Del Giudice, Nouhad Benlasfer, Philipp Mertins, Markus Landthaler, Niels H. Gehring

**Affiliations:** Institute for Genetics, University of Cologne, 50674 Cologne, Germany; Center for Molecular Medicine Cologne (CMMC), University of Cologne, 50937 Cologne, Germany; Max Delbrück Center for Molecular Medicine in the Helmholtz Association (MDC), Berlin Institute for Medical Systems Biology, Berlin, Germany; Max-Delbrück-Center for Molecular Medicine, Berlin, Germany; Cologne Center for Genomics (CCG), Medical Faculty, University of Cologne, 50931 Cologne, Germany; Berlin Institute of Health @Charite, Berlin, Germany; Institute of Biology, Humboldt-Universität zu Berlin, Berlin, Germany

**Keywords:** Nonsense-mediated mRNA decay, NMD, mRNA degradation, UPF1, gene expression, translation

## Abstract

The helicase UPF1 acts as the central essential factor in human nonsense-mediated mRNA decay (NMD) and is involved in various other mRNA degradation processes. Given its multifunctionality, distinguishing between mRNAs regulated directly and indirectly by UPF1 remains a critical challenge. We engineered two different conditional degron tags into endogenous UPF1 in human cell lines to probe the consequences of UPF1 rapid depletion. UPF1 degradation inhibits NMD within hours and strongly stabilizes endogenous NMD substrates, which can be classified into different groups based on their expression kinetics. Extended UPF1 depletion results in massive transcript and isoform alterations, partially driven by secondary effects. We define a high-confidence UPF1-regulated core set of transcripts, which consists mostly of NMD substrates. NMD-regulated genes are involved in brain development and the integrated stress response, among other biological processes. In summary, UPF1 degron systems rapidly inhibit NMD, providing valuable insights into its roles across various experimental systems.

## Introduction

In eukaryotic cells quality control mechanisms continuously monitor the transcriptome to maintain cellular integrity and prevent the expression of harmful gene products ^1^. Translation-coupled decay pathways are of specific importance among these surveillance mechanisms ^2^. They operate at the final stage of the gene expression cascade, enabling the examination and removal of transcripts that have successfully escaped previous checkpoints. One fundamental translation-coupled mechanism is nonsense-mediated decay (NMD), a process that targets mRNAs harboring premature termination codons (PTC) ^3^. PTCs introduced for example by mutations usually occur within the coding sequence and halt protein synthesis prematurely. This results in the production of C-terminally truncated proteins that are potentially dysfunctional or can even exhibit toxic properties ^4,5^. NMD serves as a crucial surveillance mechanism, ensuring that mRNAs containing PTCs are promptly eliminated to prevent the accumulation of these detrimental proteins ^6^. In mammalian cells, PTCs are identified with the help of the exon junction complex (EJC). EJCs are multi-protein complexes formed at exon-exon junctions during pre-mRNA splicing, acting as molecular markers that demarcate exon boundaries ^7–9^. When a ribosome encounters a stop codon situated at least 50 nucleotides upstream of the last exon-exon junction, the downstream EJC functions as an NMD-activating signal, triggering the recruitment of NMD factors and initiating the assembly of NMD complexes ^10–12^. This so-called 50-nucleotide rule is currently employed to annotate transcript isoforms as biotype “nonsense-mediated decay” in reference databases such as Ensembl and GENCODE^13,14^. The EJC’s capacity to activate NMD and its intrinsic characteristics explain the tendency for regular stop codons to be predominantly located in the final exon. The widespread use of alternative splicing by mammalian cells, while being crucial for expanding the coding potential of the genome ^15,16^, concurrently becomes a significant source of NMD substrates ^17,18^. This is a result of the increased probability of alternative splicing to incorporate PTC-containing sequences into internal exonic sequences. Therefore, alternatively spliced mRNAs constitute a large portion of the so-called endogenous NMD substrates expressed by every cell ^19,20^. Other NMD substrates include mRNAs with long 3′ UTRs, mRNAs with upstream ORFs, and mRNAs with 3′ UTR introns ^21^. Additionally, NMD substrates can originate from mutations, which may be the cause of genetic disorders ^22^. While in many cases NMD is triggered by the presence of an EJC, it is not required for long 3′ UTR NMD ^23,24^.

All types of NMD are tightly connected to the mRNA translation process, during which premature termination codons (PTCs) are recognized ^25^. Initially, the eukaryotic release factors (ERFs) 1 and 3 bind to the ribosome upon its stalling at a termination codon ^26^. Furthermore, ERF3 engages with the NMD machinery during translation termination either through UPF1, UPF2, or UPF3B. The details of this interaction, including whether it is direct or indirect and involves one or more of these factors, are not fully clarified at present ^27–29^. The goal of these interactions is to recruit and/or immobilize the central NMD factor UPF1 in the vicinity of the termination codon on the mRNA. Subsequently, the kinase SMG1 phosphorylates multiple SQ or TQ motifs in the unstructured regions of UPF1, found at both its N- and C-termini ^30^. Phosphorylated UPF1 is then bound by a heterodimer of SMG5 and SMG7, enabling SMG6 to endonucleolytically cleave the mRNA in close proximity to the PTC^19,31–35^. The RNA-dependent helicase UPF1 functions not only as a key component of the NMD machinery, but also participates in other mRNA degradation pathways ^36,37^. These pathways include, but are not limited to, Staufen (STAU)-mediated mRNA decay (SMD), replication-dependent histone mRNA decay (HMD), glucocorticoid-induced mRNA decay (GMD), and structure-mediated mRNA decay (SRD) ^38–41^. The common denominator of these degradation pathways is the participation of UPF1. UPF1 is recruited to the target mRNA transcripts through various RNA-binding proteins, and its participation is crucial for the effective degradation process ^36,37^. Apart from SMD, for which biochemical insights exist ^42^, details about the exact function of UPF1 remain unknown in the other cases. The NMD core machinery, including UPF1, is required for proper embryogenesis and development of many model organisms ^43–48^. A meta-analysis identified UPF1 in a core set of 188 common essential genes in mice, human cell culture and the human population ^49^. Moreover, based on current data from the Cancer Dependency Map (DepMap; 23Q2 score; https://depmap.org/portal/), UPF1 is classified as “common essential” in more than thousand human cancer cell lines ^50–55^. As a central hub in various degradation pathways, UPF1 has been extensively studied in the past. However, because UPF1 is an essential factor in mammalian cells, experimental approaches to examine its function are considerably limited. In this manuscript we employ auxin-degron mediated depletion of UPF1 in cultured human cells to characterize the transcriptome regulated by UPF1. UPF1 depletion leads to the inhibition of NMD and upregulation of known and new NMD-regulated mRNAs within a few hours. We employ combinations of various NMD-inhibited conditions to define a core UPF1-regulated transcriptome, comprising non-coding RNAs harboring small RNAs as well as mRNAs from genes implicated in brain development, RNA processing, the integrated stress response, and other biological processes. As such, the UPF1 degron system offers unparalleled and time-resolved insights into the target transcripts and functions of NMD in human cells.

## Results

The multi-domain RNA helicase UPF1 constantly monitors the transcriptome by non-specifically binding to mRNAs ^56–58^, dissociating from non-target transcripts ^59^, and initiating degradation on selected target RNAs via multiple proposed pathways such as NMD (Figures 1A-B). The loss-of-function of UPF1, as well as several other NMD and EJC factors is not tolerated in both cancer cells and the human population (Figure 1C) ^52,60^. Hence, studying the function of UPF1 poses a challenge and attempts to fully and stably delete UPF1, e.g. by conventional CRISPR/Cas9 knockout (KO), will result in lethality. Therefore, common practice involves knockdown (KD) methods based on RNA interference to transiently decrease UPF1 expression (Figure 1D) which represents one of the established methods to inactivate NMD. However, efficiently downregulating UPF1 in various cell types typically results in reduced proliferation, cell cycle arrest, or apoptosis, which impedes long term KD experiments ^61–64^. Here, we assess UPF1-KD as an experimental approach to inhibit NMD (Figure 1D) compared to combined SMG6 and SMG7 downregulation, which strongly inhibits NMD activity ^19,65,66^. We downregulated UPF1 in human embryonic kidney cells (HEK293) and the colon cancer cell line HCT116 for 48-72 hours by using either an established siRNA sequence ^67^ or a high complexity pool of 30 siRNAs (siPOOL) to reduce off-target effects ^68^ (Figures 1E and S1A). A well-characterized alternatively spliced NMD target SRSF2^69^ and two NMD-sensitive lncRNAs, ZFAS1 and GAS5^70^, showed stronger accumulation in SMG6+SMG7 KD compared to both UPF1 KD conditions (Figures 1F-G and S1B-C). Despite strongly reduced UPF1 protein levels, residual UPF1 appears to sustain partial NMD activity, particularly in early timepoints. This aligns with mass spectrometry-based data from HeLa cells ^71^, 29 human tissues ^72^, or mouse fibroblasts ^73^, which determined high UPF1 abundance (> 250,000 copies per cell) and calculated long UPF1 protein half-life (T1/2 ≥48h) (Figures S1D-G). In contrast, both SMG6 and SMG7 are about 100 times less abundant and turn over more rapidly, explaining the faster onset of NMD inhibitory effects following their KD. To obtain a global view on NMD-inhibition, we investigated the transcriptome-wide effects of UPF1 KD across different human cell lines by analyzing recently published RNA-Seq datasets ^74–80^. This strategy provides a broader perspective on UPF1 KD and aims to mitigate individual biases, since many of these datasets were generated in different laboratories using different KD protocols, target sequences, and cell lines. For comparison, we re-analyzed RNA-Seq data from HEK293 cells subjected to either combined SMG7-KO + SMG6-KD or the double-KO of both UPF3 paralogs (UPF3A & UPF3B) ^19,81^. Differential gene expression (DGE) analysis of significantly regulated genes, as well as differential transcript expression (DTE) analysis of NMD-annotated transcripts, confirmed the strong transcriptome-wide effects and upregulation of NMD targets in cells depleted of SMG6/7 or UPF3A/B (Figure 1H and S1H). These observations are also supported by the increased levels of individual, previously identified NMD-sensitive genes/isoforms, such as GADD45B or SRSF2^69,82^. In contrast, among the eight datasets examined, only the UPF1-KD dataset with the strongest reduction of UPF1 mRNA levels ^79^ showed comparable NMD inhibitory effects. Technical characteristics of the RNA-Seq data, such as sequencing-depth, mapped reads, or library preparation, might contribute to the apparently weaker effects of UPF1-KD on NMD in some datasets (Figure S1I), although overall the accumulation of NMD-annotated transcripts correlates well with the extent of UPF1 mRNA reduction (Figure S1J). In conclusion, the transient downregulation of UPF1 via RNAi results in only partial NMD inhibition compared to more potent NMD-compromising conditions.

**Figure 1.**
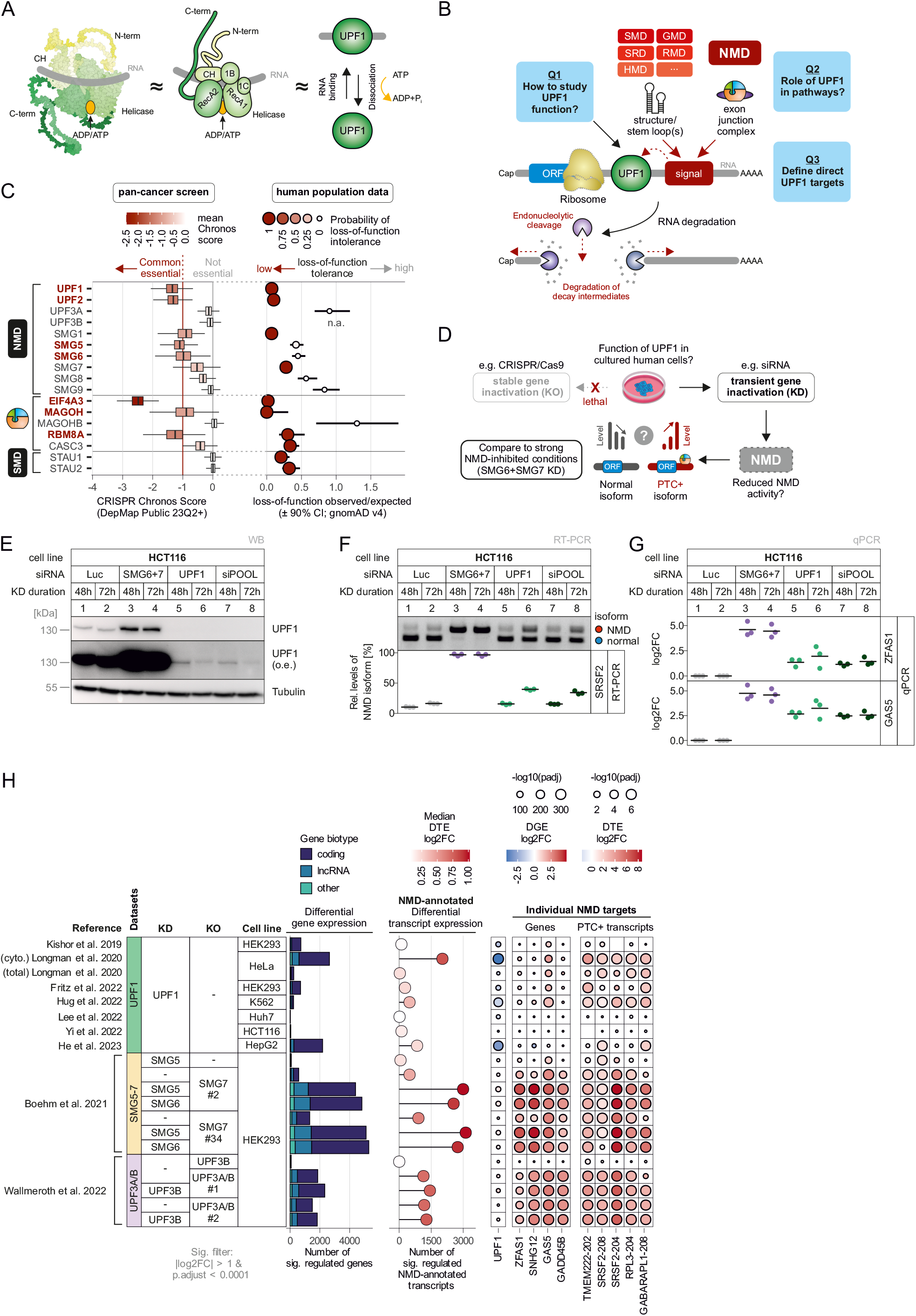
Knockdown of the essential RNA-binding protein UPF1 results in incomplete NMD inhibition. (**A**) Schematic representation of UPF1 RNA-binding function and protein domains based on AlphaFold prediction (AF-Q92900-F1-model_v4) and UPF1-RNA-ADP:AlF4 crystal structure (PDB ID 2xzo). (**B**) Schematic overview of UPF1 function in RNA degradation pathways, such as nonsense-mediated decay (NMD), Staufen-mediated mRNA decay (SMD), glucocorticoid-induced mRNA decay (GMD), structure-mediated mRNA decay (SRD), Regnase-mediated mRNA decay (RMD) or replication-dependent histone mRNA decay (HMD). Signals such as stem loops or the exon junction complex in the 3′ UTR of actively translated mRNAs lead to UPF1-mediated degradation. The key questions of this study are highlighted. (**C**) Gene essentiality scores from a pan-cancer screen (DepMap) or loss-of-function observed/expected ratios (± 90% confidence interval; CI) from human population data (gnomAD v4) are shown for individual NMD, EJC and SMD factors. The Chronos score is depicted as boxplot, the box extends to the 25th and 75th percentile with the median in bold line, outliers are not shown. (**D**) Schematic overview of the workflow to study UPF1 function in NMD in human cells. (**E**) Western blot showing levels of UPF1 protein after knockdown in HCT116 cells with different siRNAs for 48 or 72h, Tubulin serves as loading control, o.e. = overexposure. (**F**) Detection of the NMD target SRSF2 by end-point RT-PCR in the indicated knockdown conditions. The NMD and normal isoforms are indicated and the relative mRNA levels of SRSF2 isoforms were quantified from the agarose gel bands, individual data points and mean are shown (n=3). (**G**) Probe-based qPCR of NMD targets ZFAS1 and GAS5, shown as log2 fold change (log2FC), individual data points and mean are shown (n=3). (**H**) Comparison of various published RNA-Seq data regarding the number of significantly regulated genes (padj < 0.0001 & abs(log2FC) > 1) stratified by GENCODE biotype (left), the number and median log2FC of significantly regulated GENCODE NMD-annotated transcripts (middle), as well as expression changes of UPF1 mRNA or individual NMD target genes and transcripts (right).

### Characterizing an AID-degron system for rapid and functional UPF1 depletion

Given the lethality of genomic UPF1 KO and the moderate effects obtained with UPF1 KDs, we aimed to explore the potential for selectively depleting UPF1 at the protein level. We employed the auxin-inducible degron (AID) technology to conditionally degrade UPF1 protein within the cellular environment ^83^. To this end, we inserted either an N-terminal or C-terminal V5-AID-tag (AID = miniIAA7 = AtIAA7 amino acids 37–104) into the genomic UPF1 locus of HCT116 cells, which contained the auxin receptor F-box protein-encoding AtAFB2-mCherry in the AAVS1 locus ^84^ (Figure 2A and Table S1). As an initial functional test, we performed shallow RNA-Seq (≈22 × 10^6^ reads per sample) of control (untagged UPF1) as well as N-terminally and C-terminally AID-tagged UPF1 cell lines treated with 500 µM indole-3-acetic acid (IAA) for various timepoints (0-12h) (Figure S2A). Both AID-tagged cell lines showed a time-dependent accumulation of NMD-annotated mRNAs during IAA incubation, whereas the addition of IAA did not result in substantial changes in gene expression in control cells. Based on our initial RNA-Seq results, we chose the N-terminally AID-tagged UPF1 cell line for subsequent experiments. Proteomic analyses revealed that IAA treatment for 6-24h strongly and specifically reduced UPF1 protein levels (Figure S2B), whereas other potential UPF1 interactors (Figure S2C) or NMD/EJC/SMD factors are largely unaffected or upregulated (Figure 2B). Therefore, the UPF1 degron cells enable the specific and rapid depletion of UPF1 protein with minimal side effects of the IAA treatment, allowing a time-resolved investigation of NMD inhibition. We conducted an extended time course of IAA treatment, spanning up to 48h and including an early 2-hour timepoint (Figure S2D). Western blot analysis confirmed the rapid and strong depletion of UPF1 protein even after 2h IAA incubation (Figure 2C). Selected mRNAs showed accumulation of NMD-sensitive transcripts over time, exhibiting seemingly different kinetics. The alternatively spliced NMD-sensitive SRSF2 isoform reached maximal relative levels after 12 hours, whereas the snoRNA host genes ZFAS1 and GAS5 continued to accumulate up to 48h (Figure 2C). These different accumulation patterns could be influenced by transcription and processing kinetics, translation rates and/or NMD efficiency. To test this hypothesis, we performed deep RNA-Seq of the extended time course samples (*≈*81 × 10^6^ reads per sample). The analysis of NMD-annotated transcripts and individual RNA targets revealed a pronounced time-dependent NMD-inhibition (Figure 2D). Within four to eight hours of UPF1 depletion, NMD inhibition surpassed SMG7 + SMG6 depletion or the strongest published UPF1 KD dataset (Figure 2D). Of note, we found no substantial expression changes of UPF1 mRNA in the IAA-treated AID-UPF1 cells, indicating that the UPF1 depletion occurs exclusively on the protein level. Consistent with the PCR results, visual inspection of SRSF2 mapped reads confirmed the complete isoform switch from the canonical isoform to the NMD-annotated transcripts within 12h of UPF1 depletion (Figure 2E). Principal component analysis of global gene-level counts revealed a nearly linear trend along principal component 1 from 0h to 8h/12h, whereas further IAA incubation time up to 48h spans mostly along principal component 2 (Figure 2F). One possible explanation for this observation is that direct effects of NMD inhibition manifest in the first 8h to 12h, whereas additional (secondary) effects predominantly emerge after prolonged UPF1 depletion.

**Figure 2.**
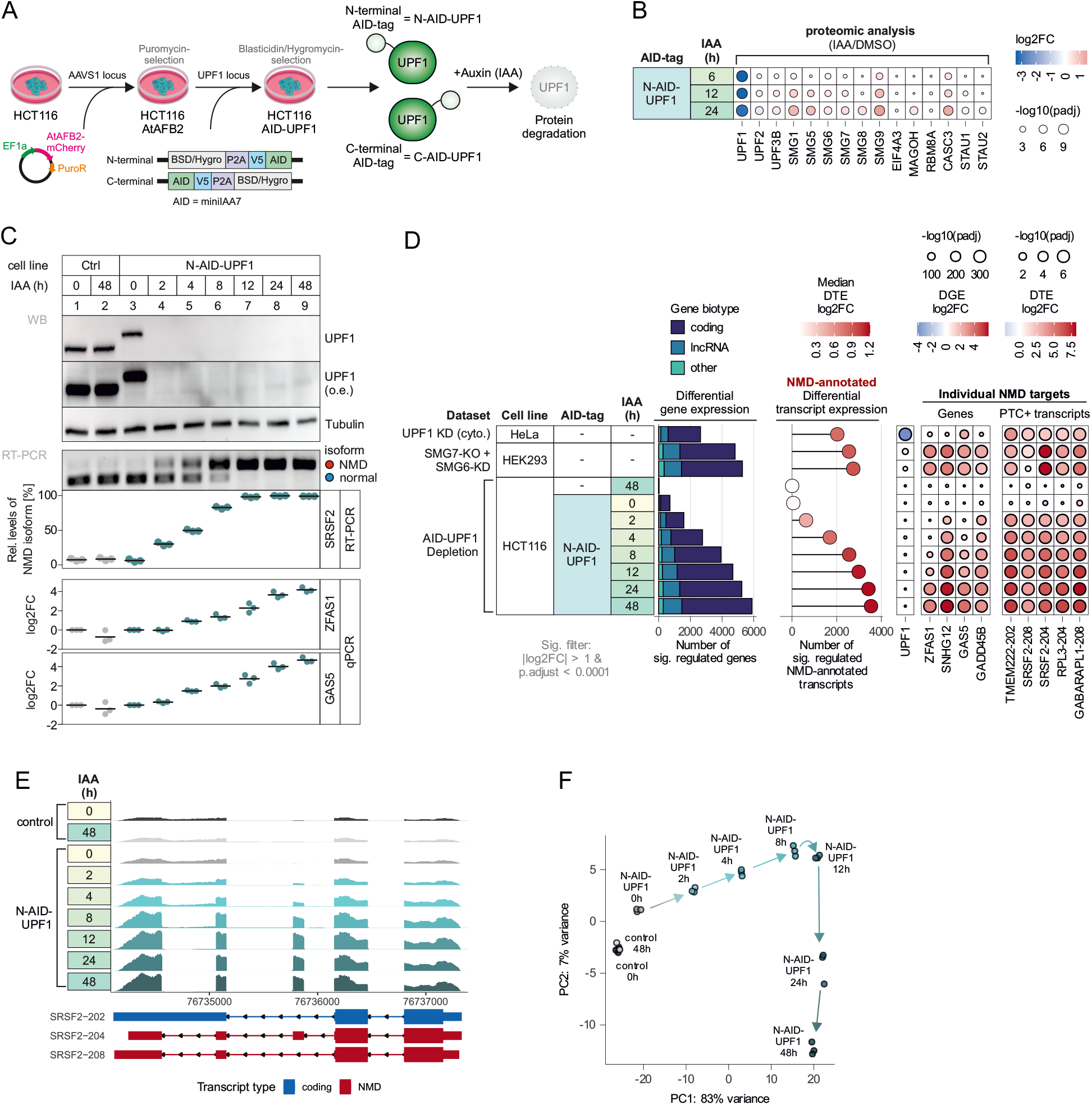
Rapid depletion of UPF1 via the auxin-inducible degron (AID) system inhibits NMD. (**A**) Schematic representation of cell line generation allowing the auxin-inducible (indole-3-acetic acid; IAA) depletion of UPF1. For details see Methods. (**B**) Protein expression levels of NMD/EJC/SMD factors in IAA-treated versus DMSO control conditions, determined by mass spectrometry. (**C**) (Top) Western blot showing levels of UPF1 protein in control (Ctrl) or N-terminal AID-tagged UPF1 HCT116 cell lines, treated with 500 µM IAA for the indicated time. Tubulin serves as loading control, o.e. = overexposure. (Middle) Detection of the NMD target SRSF2 by end-point RT-PCR. The relative mRNA levels of SRSF2 isoforms were quantified from the agarose gel bands, individual data points and mean are shown (n=3). (Bottom) Probe-based qPCR of NMD targets ZFAS1 and GAS5, shown as log2 fold change (log2FC), individual data points and mean are shown (n=3). (**D**) Comparison of N-AID-UPF1 with SMG7-KO + SMG6-KD (Boehm et al. 2021) or cytoplasmic UPF1 KD (Longman et al. 2020) RNA-Seq data regarding the number of significantly regulated genes (padj < 0.0001 & abs(log2FC) > 1) stratified by GENCODE biotype (left), the number and median log2FC of significantly regulated GENCODE NMD-annotated transcripts (middle), as well as expression changes of UPF1 mRNA or individual NMD target genes and transcripts (right). (**E**) Read coverage of SRSF2 gene from N-AID-UPF1 RNA-Seq data. Y-axis (reads) was scaled equally for all conditions and biologically relevant annotated transcripts are shown below. (**F**) Principal component analysis of gene-level counts from N-AID-UPF1 RNA-Seq data, arrows were added to visualize the time course.

We first approached the above-mentioned hypotheses on the gene-level by clustering significantly up- or down-regulated genes according to their time-dependent normalized counts. We included genes that were significantly regulated at least at one timepoint compared to control condition (untagged UPF1 with 0h IAA; Table S2). For both up- and down-regulated genes, five clusters were identified that we described according to their expression profiles as “early”, “delayed”, “late”, “inverse” and “peak” (Figures 3A and S3A-B). The early and peak clusters reached plateaus or peak amplitudes between only 8-12h of IAA treatment, suggesting that these gene expression changes are a direct consequence of UPF1 depletion. In contrast, the expression changes in the late cluster could be only observed after 24h-48h of IAA addition and therefore likely represent secondary effects. We hypothesize that, similar to the early clusters, the delayed clusters primarily result from the loss of UPF1 protein. However, these genes have either slower upregulation kinetics or a greater accumulation capacity. The inverse clusters are already mis-regulated at 0h IAA treatment and likely artifacts of the AID tagging. Of all GENCODE-annotated genes, we found more than two-thirds of protein-coding genes to be sufficiently expressed to allow expression analyses, whereas most lncRNAs or other biotype genes had too few RNA-Seq counts (Figure 3B). Furthermore, a substantial proportion of protein-coding genes were detected to be differentially expressed in the time course experiment, suggesting that UPF1 primarily regulates transcripts that are usually translated (Figures 3B-C).

**Figure 3.**
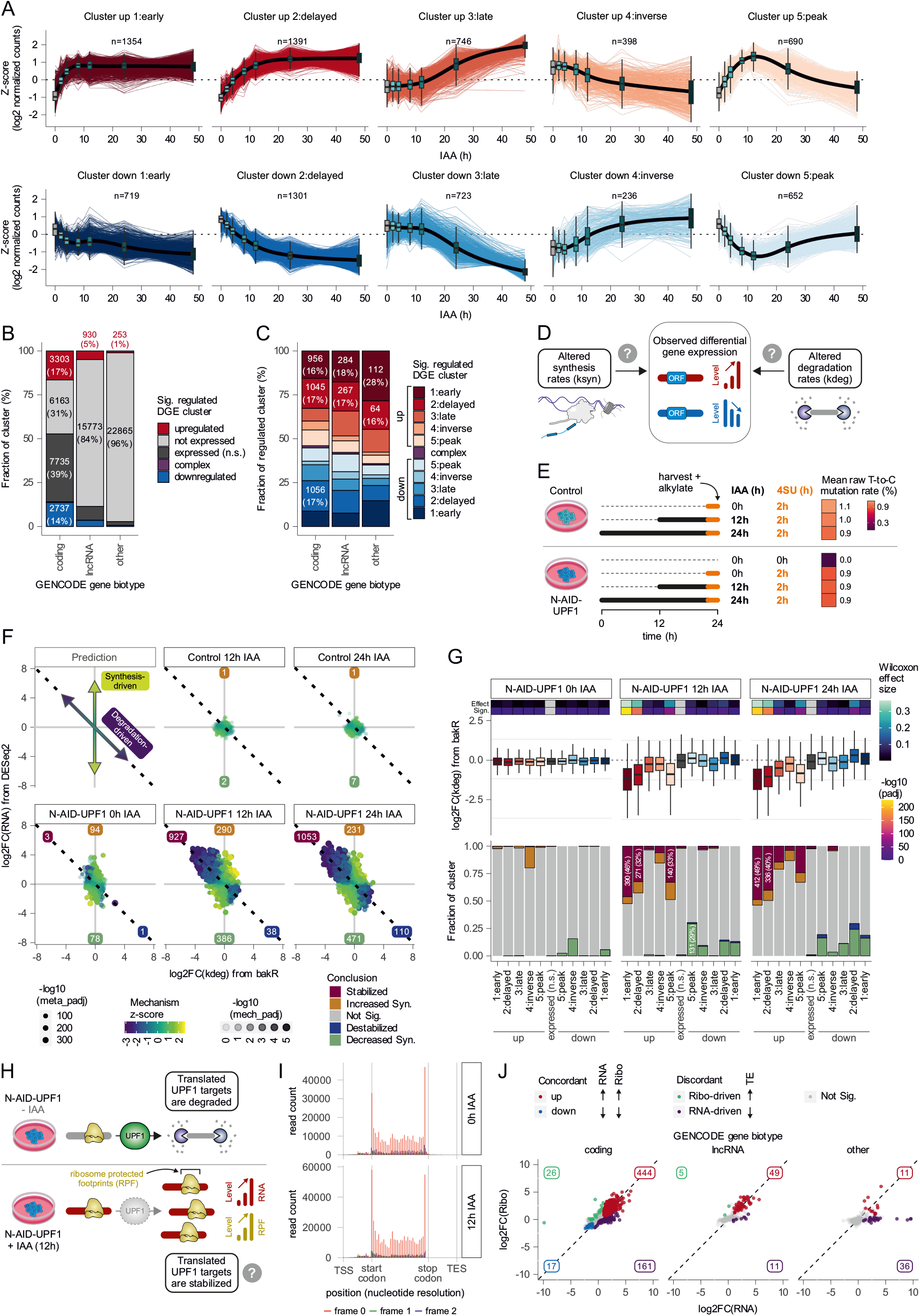
UPF1 depletion allows the identification of upregulated, stabilized and translated target genes. (**A**) Clustering of N-AID-UPF1 RNA-Seq data based on Z-scaled log2-transformed gene-level counts (k-means, k = 5) of upregulated (top row) or downregulated (bottom row) genes is depicted as Z-score over IAA treatment in hours. Genes were selected as up- or downregulated when at least one timepoint was significant compared to control 0h IAA condition (padj < 0.0001 & abs(log2FC) > 1). Number of genes per cluster are indicated. (**B**) Barplot showing the proportions of GENCODE-annotated genes per indicated biotype classified as up- or downregulated (= identified in clusters in (A)), complex (= identified in both up- and downregulated clusters), expressed without significant changes (n.s. = not significant) or found as not expressed (< 10 counts per half of conditions) in the N-AID-UPF1 RNA-Seq data. (**C**) As in (B) proportions of genes per GENCODE biotype identified in individual significantly regulated differential gene expression (DGE) clusters are shown. (**D**) Scheme depicting the potential mechanistic explanation for gene expression changes. (**E**) Schematic overview of experimental setup for SLAM-seq metabolic labelling of control or N-AID-UPF1 HCT116 cells with 4-thiouridine (4SU). The mean raw T-to-C mutation rate as determined by bakR is shown per condition. (**F**) Mechanistic dissection of SLAM-seq data by integrating the detected changes in RNA levels (log2FC(RNA) by DESeq2) and degradation rate (log2(kdeg) by bakR) in each condition compared to control 0h IAA HCT116 cells. Size of points indicates the combined statistical significance of bakR and DESeq2 analysis (meta_padj), the mechanism score Z-score (fill color) indicates degradation-driven (negative values) or synthesis-driven (positive values) expression changes and transparency of points depicts the statistical significance of mechanism scores (mech_padj). Conclusions are based on parameters defined in the Methods section and the numbers of genes for each conclusion type are indicated. (**G**) (Top) Boxplot of degradation rate changes in the indicated conditions of the N-AID-UPF1 SLAM-seq data stratified by DGE clusters defined in (A). The Wilcoxon effect size was calculated and the statistical significance was determined using two-sided Dunn’s test of multiple comparisons with the not significant (n.s.) expressed cluster as reference. (Bottom) Barplot showing the proportion of genes of each DGE cluster classified by the conclusion determined in (F). (**H**) Schematic overview of translation-dependent degradation of UPF1 target genes and postulated detection of increased ribosome protected footprints for target genes upon UPF1 depletion. (**I**) Metagene plot of P-site read coverage aggregating signal over all covered transcripts. (**J**) Combined analysis of RNA expression changes (log2FC(RNA)) and ribosome occupancy changes (log2FC(Ribo)) of 12h versus 0h IAA condition, stratified by GENCODE biotype. Details for classification of genes in concordant or discordant classes are defined in the Methods section. The numbers of genes for each class are indicated.

### Transcriptome changes in UPF1-depleted cells result from mRNA stabilization

We initially considered whether the inherent transcript half-lives could explain the clustering results. However, when comparing individual clusters with published mRNA half-lives from HEK293 cells ^85^, no apparent dependence was found (Figure S3C). Next, we explored whether the changes in RNA expression levels in UPF1-depleted cells result from altered synthesis (ksyn) or degradation (kdeg) rates (Figure 3D). To this end, we performed SLAM-seq of control and AID-UPF1 cells at three timepoints of IAA treatment (0, 12 and 24 hours) with 4-thiouridine (4SU) labelling for 2h prior to harvesting (Figure 3E) ^86^. T-to-C conversion rates were around 1% for all labeled samples and therefore usable for downstream analyses via bakR (Figure 3E) ^87,88^. By calculating log2-fold changes in the degradation rate constant [log2FC(kdeg)] with differential RNA expression level analysis from DESeq2 [log2FC(RNA)], we attribute gene expression regulation to synthesis-driven and/or degradation-driven processes (Figure 3F). Genes with altered RNA levels but unchanged degradation rate constants are synthesis-driven (positive mechanism z-score; along y-axis), whereas degradation-driven genes exhibit coordinated changes in both RNA and kdeg (negative mechanism z-score; along diagonal dashed line). According to this classification, UPF1 depletion resulted in stabilized transcripts from 927 or 1053 genes after 12 hours or 24 hours of IAA treatment, respectively (Figure 3F and Table S3). Of note, IAA treatment for 12 or 24 hours also resulted in about 700 genes with synthesis-driven expression changes, suggesting that transcriptional responses contribute to the altered transcriptome in UPF1-depleted cells. Gene ontology analyses of these synthesis-driven genes in the 12-hour timepoint revealed e.g. reduced expression of energy/respiration-related genes and increased expression of stress-, apoptosis-, and metabolism-related genes (Figure S3D). Combining the results of the SLAM-seq experiment with the time course DGE clustering showed that the upregulated early, delayed, and peak clusters have a strong and significant tendency for reduced degradation rates [log2FC(kdeg)] and being classified as stabilized (Figure 3G). In conclusion, efficiently depleting UPF1 over time revealed distinct clusters of transcripts exhibiting varying kinetics of differential expression. The increased expression levels in the early and delayed upregulated clusters are primarily caused by a reduction in RNA degradation rates.

### UPF1-targeted mRNAs undergo translation

Considering that a large portion of UPF1-dependent mRNA degradation pathways depend on active translation (reviewed in ^37^), we next examined the translatome of UPF1-depleted cells. In these cells, direct targets of UPF1 are anticipated to be stabilized at the RNA level, accompanied by a proportional increase in ribosome occupancy (Figure 3H). Analysis with Ribo-seQC ^89^ showed that more than half of the reads mapped to annotated CDS (Figures S3E-F) with an overall reading frame preference of around 75% and a good 3-nucleotide periodicity (Figures 3I and S33G). We reliably detected Ribo-Seq reads for 40-70% of the genes, depending on the DGE cluster type (Figure S33H). The majority of genes with significant changes were concordantly up-regulated in Ribo-Seq and mRNA-Seq (Figure 3J). This was not only the case for annotated protein-coding genes, but also for 49 lncRNAs including ZFAS1 and GAS5, non-coding RNAs with expressed mini ORFs^90^. Of note, we only detected few genes with concordantly downregulated RNA and ribosome occupancy levels. UPF1 depletion also caused discordant changes, e.g. increased ribosome occupancy on genes with unchanged RNA levels. Analysis of translation efficiency (TE) ^91^ identified around 200 genes with unchanged ribosome, but increased RNA levels (Figure 3J and Table S4). This result indicates that not all UPF1-dependent upregulated genes result solely from translation-mediated processes.

### Full reversibility of UPF1 depletion provides insights into the recovery of the NMD-targeted transcriptome

To further dissect the role of UPF1 in shaping the transcriptome, we utilized the reversibility of the AID system ^83^. After successful depletion of UPF1 with auxin, transferring the cells to medium without auxin allows UPF1 protein repletion. This enables studying how transcripts/genes are targeted by the recovering UPF1 protein expression in an NMD-compromised background. We initially treated N-AID-UPF1 cells for 24h with IAA, removed the auxin and harvested cells at various time-points up to 48h without IAA (Figure S2D). Western blot analysis showed increasing UPF1 protein abundance reaching control levels after 24-48h of recovery (Figure 4A). Selected NMD targets strongly accumulated after 24h of UPF1 depletion, but their levels normalized upon UPF1 recovery (Figure 4A). Deep RNA-Seq of these samples revealed that the most striking transcriptome-wide changes occurred after 4-12 hours of UPF1 re-expression (Figure 4B). We then explored whether the gene clusters identified earlier (Figure 2) exhibit corresponding responses to both UPF1 depletion and repletion. Specifically, we expected rapid down-regulation of direct UPF1 targets upon UPF1 recovery. Indeed, the genes belonging to the early and peak clusters responded more quickly, whereas the late cluster exhibited prolonged mis-regulated gene expression (Figure 4C). To explore the time-dependent gene expression changes in more detail, we employed ImpulseDE2 to fit interpretable models to the RNA-Seq counts of each gene in the UPF1 depletion, recovery, or the combination of both datasets (Figure S4A) ^92^. For more than 10000 identified genes with significant expression changes over time the best fitting model was determined, which either captured monotonous (sigmoid model) or transient (impulse model) gene expression changes (Figure S4A and Table S5) ^93^. The majority of detected genes in the UPF1 depletion or recovery dataset showed sigmoidal expression patterns, reflecting their monotonous up-/downregulation over time. In contrast, most genes in the combined depletion/recovery time course followed an impulse pattern, which confirms the reversibility of expression changes (Figure S4A). We extracted informative onset (t1) and offset (t2) times, which are the transition times from initial to peak state (t1) and peak to steady state (t2). These emphasize the predominantly fast expression response after UPF1 depletion or recovery (Figures S4A-B). Next, we used the model parameters to reevaluate gene classification into DGE cluster using the combined depletion/recovery data (Figure S4C). Onset times aligned closely with our previous categorization, especially for the early 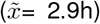, delayed 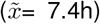 and late 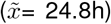 upregulated clusters (Figure 4D). Interestingly, the delayed and early clusters exhibited similar short offset times, suggesting that re-expressed UPF1 targets both clusters equally for rapid degradation. Overall, these observations substantiate our previous statements that genes in the early and delayed cluster are directly targeted by UPF1.

**Figure 4.**
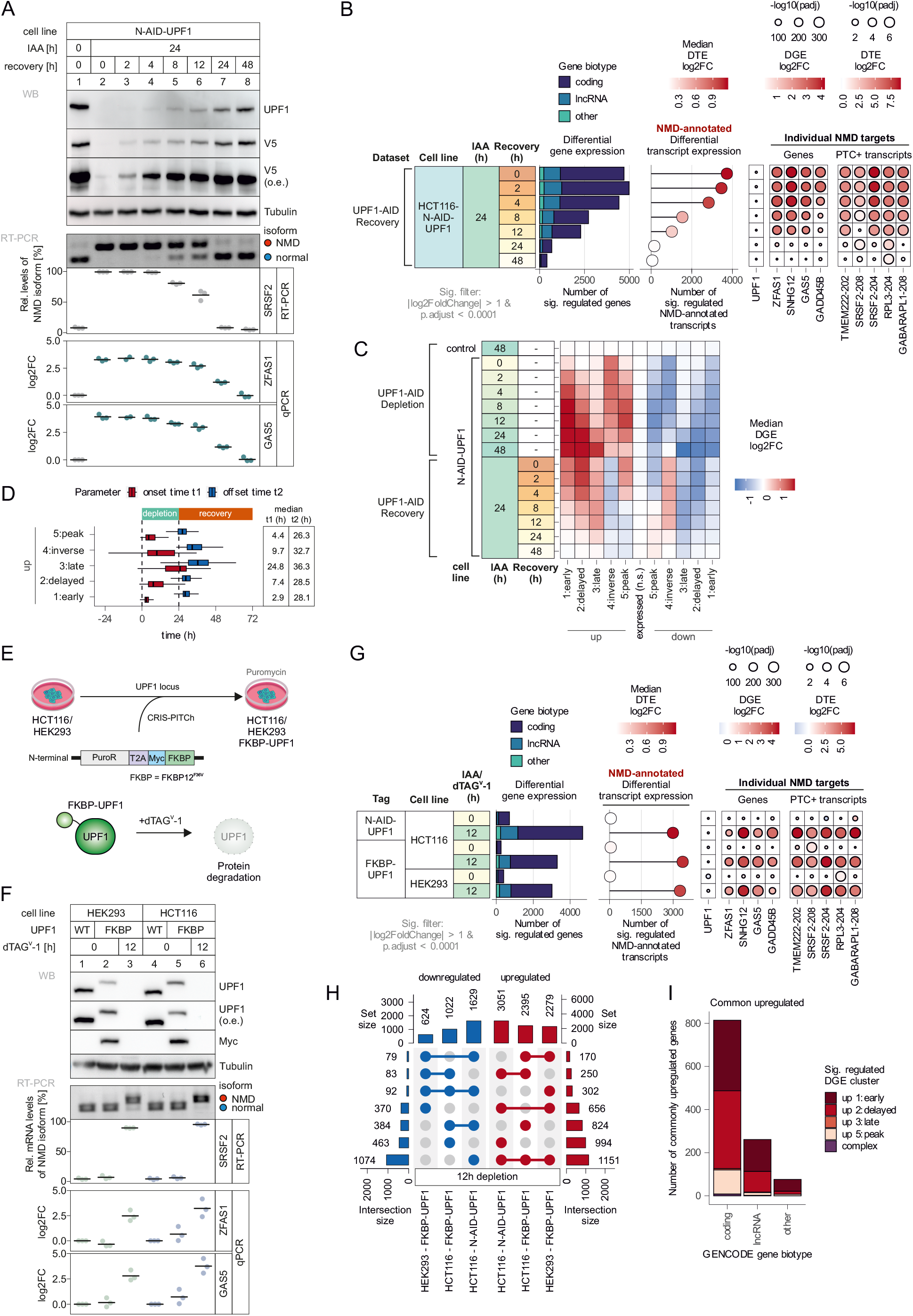
Recovery of UPF1 depletion and FKBP-based degron systems provide robust identification of UPF1 targets. (**A**) (Top) Western blot showing levels of UPF1 protein in N-terminal AID-tagged UPF1 HCT116 cell line treated with 500 µM IAA for the indicated time and incubated for the recovery time without auxin. V5-AID-tagged UPF1 was also detected via V5 antibodies. Tubulin serves as loading control, o.e. = overexposure. (Middle) Detection of the NMD target SRSF2 by end-point RT-PCR. The relative mRNA levels of SRSF2 isoforms were quantified from the agarose gel bands, individual data points and mean are shown (n=3). (Bottom) Probe-based qPCR of NMD targets ZFAS1 and GAS5, shown as log2 fold change (log2FC), individual data points and mean are shown (n=3). (**B**) RNA-Seq data of N-AID-UPF1 depletion/recovery regarding the number of significantly regulated genes (padj < 0.0001 & abs(log2FC) > 1) stratified by GENCODE biotype (left), the number and median log2FC of significantly regulated GENCODE NMD-annotated transcripts (middle), as well as expression changes of UPF1 mRNA or individual NMD target genes and transcripts (right). (**C**) Median gene expression changes per DGE cluster for both N-AID-UPF1 depletion and recovery RNA-Seq datasets. (**D**) ImpulseDE2-derived modelled onset (t1) and offset (t2) time parameters for the indicated DGE cluster in context to the experimental setup are depicted as boxplot. Median values are shown. (**E**) Schematic representation of cell line generation allowing the dTAGV-1-inducible depletion of UPF1. For details see Methods. (**F**) (Top) Western blot showing levels of UPF1 protein in control or N-terminal FKBP-tagged UPF1 HEK293 or HCT116 cell lines treated with 0.25 µM dTAGV-1 for the indicated time. Myc-FKBP-tagged UPF1 was also detected via Myc antibodies. Tubulin serves as loading control, o.e. = overexposure. (Middle) Detection of the NMD target SRSF2 by end-point RT-PCR. The relative mRNA levels of SRSF2 isoforms were quantified from the agarose gel bands, individual data points and mean are shown (n=3). (Bottom) Probe-based qPCR of NMD targets ZFAS1 and GAS5, shown as log2 fold change (log2FC), individual data points and mean are shown (n=3). (**G**) RNA-Seq data of N-AID-UPF1 and FKBP-UPF1 depletion regarding the number of significantly regulated genes (padj < 0.0001 & abs(log2FC) > 1) stratified by GENCODE biotype (left), the number and median log2FC of significantly regulated GENCODE NMD-annotated transcripts (middle), as well as expression changes of UPF1 mRNA or individual NMD target genes and transcripts (right). (**H**) UpSet plot of the overlap of significantly up- or downregulated genes (padj < 0.0001 & abs(log2FC) > 1) between AID- and FKBP-mediated UPF1 depletion for 12h. (**I**) Barplot showing the number of genes upregulated in all three UPF1-depletion conditions from (H) per DGE cluster and stratified by GENCODE biotype.

### NMD-regulated gene expression profiles are highly reproducible across different degron systems and cell lines

All UPF1 depletion results obtained so far were obtained with the AID system in HCT116 cells. To evaluate the robustness of UPF1 depletion, we inserted an N-terminal dTAG (FKBP12^F36V^) ^94^ into the genomic UPF1 locus in HCT116 and commonly used HEK293 cells (Figure 4E). After 12h of dTAG^V^-1 treatment, FKBP-tagged UPF1 was not detectable by Western blot analysis (Figure 4F). Concurrently, the SRSF2 NMD isoform, ZFAS1 and GAS5 strongly accumulated, suggesting a highly effective inhibition of NMD (Figure 4F). In comparison to AID-UPF1, cells with depleted FKBP-UPF1 globally showed regulation of significantly fewer genes. However, NMD inhibition was similar, indicated by the upregulation of NMD-annotated transcripts and individual target RNAs. (Figure 4G). When analyzing the overlap of up- or downregulated genes between the different degrons, we found that most upregulated genes were shared between all three UPF1-depleted conditions (Figure 4H). In contrast, only a limited number of genes were concurrently downregulated in two or three UPF1-depleted cell lines. Interestingly, a large portion of the 1151 commonly upregulated genes had previously been classified in the early or delayed upregulated clusters (Figure 4I), demonstrating the robustness of our classification even when employing a different degron system or cell line.

### The isoform-resolved UPF1-regulated transcriptome complements gene-level analysis and provides a comprehensive view of UPF1 regulation

Next, we investigated the effects of UPF1 depletion on the transcriptome by using transcript isoform-specific approaches, which allow different types of analyses (Figure S1H). First, we quantified the number of differentially expressed transcripts deemed significant in all HCT116 and HEK293 UPF1-depleted conditions, stratified by GENCODE transcript biotype annotation (Figure 5A). Consistent with the gene-level analyses, the number of regulated transcripts increased during the depletion and decreased with the recovery of UPF1. The majority of regulated transcripts were annotated as protein-coding, followed by NMD-annotated transcripts. Specifically, when UPF1 was absent, there was a distinct tendency for upregulation observed in the latter category (Figure 5A). Clustering of all differentially expressed transcripts revealed in total 25763 upregulated and 20472 downregulated mRNA isoforms distributed over 5 clusters each (Figures S5A-B). Clusters 4 and 5 primarily contained events specific to the HCT116 or HEK293 cell line, respectively, whereas the clusters 1-3 resembled the early, delayed and late clusters identified in our previous gene-level analysis (Figures 5B and S5A-B). About 28% of the transcripts annotated as NMD targets were upregulated when UPF1 was depleted and almost 70% of differentially regulated NMD isoforms were found in cluster 1 or 2 (Figure 5C-D). Conversely, one third of annotated NMD target transcripts were not detected due to low counts (Figure S5C), while another third did not change their expression significantly in our datasets (Figure 5C). Next, we investigated the overlap in differentially regulated transcripts resulting from AID- and FKBP-mediated UPF1 depletions in both HCT116 and HEK293 cells. Consistent with the gene-level analysis, we found limited overlap in downregulated isoforms, whereas the majority of upregulated isoforms (n=4477) were consistently shared across all three conditions (Figure 5E). Nearly all of the commonly shared upregulated transcripts belonged to clusters 1 (early) and 2 (delayed), with more than 1500 NMD-annotated transcripts representing the predominant biotype (Figure 5F). In contrast to the gene-level analyses, transcript-level approaches allow for the detailed examination of transcript properties such as length, GC-content or predicted structure in the 3′ UTR (Figure 5G). This analysis is particularly insightful as certain UPF1-dependent decay pathways rely, at least in part, on long, GC-rich, and/or structured 3′ UTRs^39,95^. We first determined whether there is a correlation between these transcript properties and the expression changes in any UPF1 condition. Overall, the coefficient of determination (R^2^) is very low, indicating no general dependency of the transcript expression on the tested properties (Figure 5H). We analyzed the relationship between the transcript properties and their expression clusters, excluding cell line specific clusters. Compared to the background (unchanged expressed transcripts), many significant differences were detected, however only with small effect sizes and mostly unexpected relations (Figure 5I). For example, longer 3′ UTR lengths were found in downregulated cluster 2 and lower transcript GC content in downregulated cluster 1. Interestingly, transcripts in the upregulated cluster 1 consist of significantly more exons. We also conducted an analysis of differential transcript usage (DTU; proportional differences in the transcript composition of one gene; Figure 5J) and observed comparable results to those obtained from the differential transcript expression (DTE) analysis (Figure 5A). Combining transcript-specific analyses (DTE and DTU) with differential gene expression (DGE) on the gene-level, we find that with increasing duration of UPF1 depletion, up to 65% of the genes undergo at least one type of transcriptome perturbation (Figure S5D). Furthermore, we employed LeafCutter to detect alternative splicing in an annotation-free manner ^96^. This revealed that thousands of significant intron clusters contain at least one unannotated junction in UPF1-depleted cells (Figure 5J). Interestingly, the cryptic alternative splicing changes increased with prolonged UPF1 depletion. One class of relevant target mRNAs are the serine/arginine-rich splicing factor (SR) family genes, which encode for regulators of alternative and constitutive splicing (reviewed in ^97^). SR gene expression is (auto-)regulated by unproductive splicing coupled to degradation via NMD, e.g. by poison exon inclusion ^98–100^. Indeed, we observed for most SR genes strong decrease of the protein coding isoforms and increase of the NMD-annotated isoforms upon UPF1 depletion (Figure 5J). Therefore, inhibition of NMD causes the rapid mRNA isoform mis-regulation of splicing regulators like SR proteins, which could contribute to the observed increasing usage of cryptic splice sites. As SR protein half-lives were predicted to be rather long (mean T1/2 > 60 h; Figure S5E), this effect will probably manifest especially during long-term inactivation of NMD.

**Figure 5.**
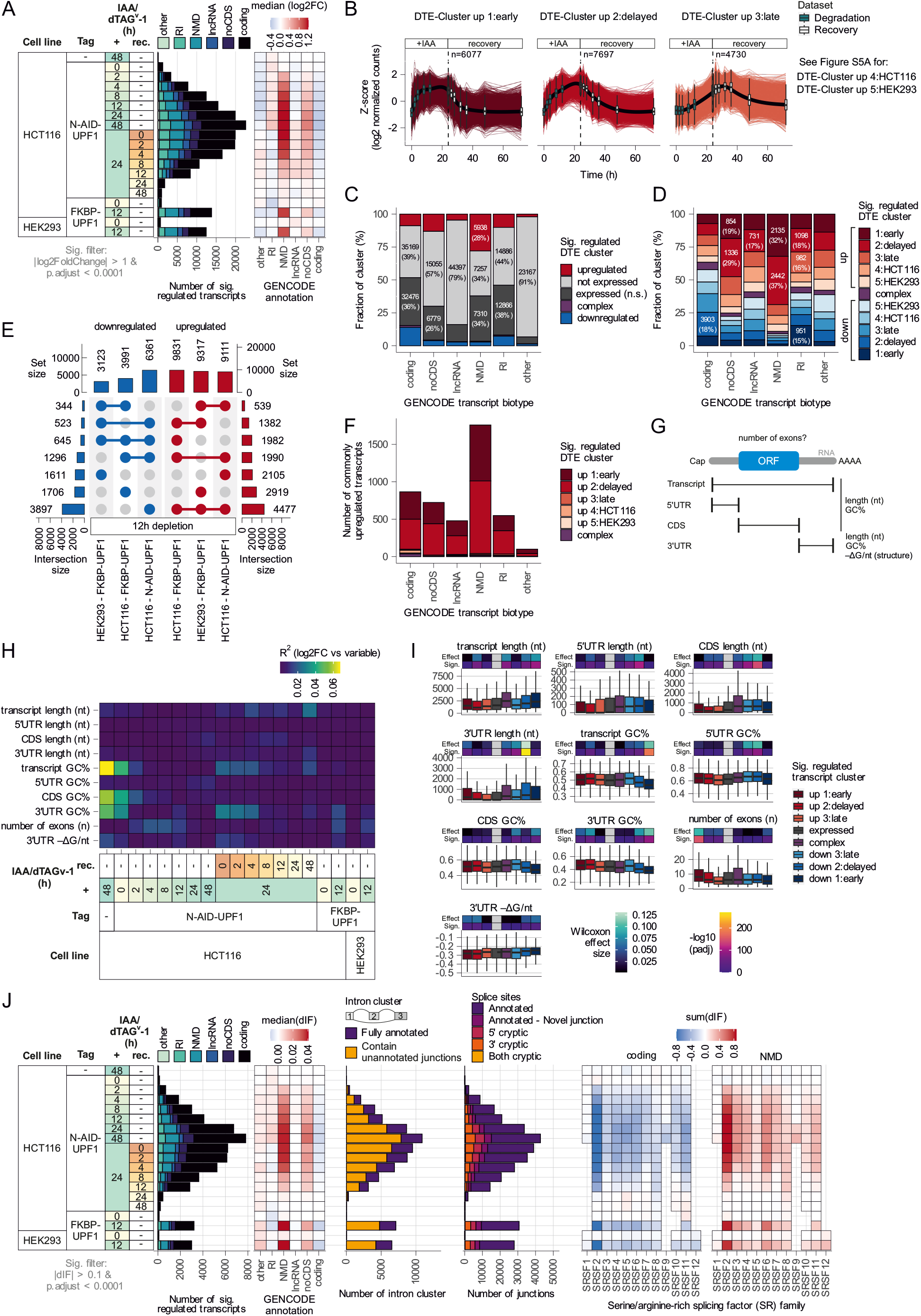
UPF1 depletion allows in-depth characterization of targeted mRNA isoforms. (**A**) Differential transcript expression (DTE) analysis of AID- and FKBP-mediated UPF1 depletion/recovery RNA-Seq datasets, showing the number of significantly regulated mRNA isoforms as well as median expression changes per GENCODE transcript biotype. (**B**) Z-scaled log2-transformed transcript-level counts of upregulated transcripts per identified DTE cluster for both depletion and recovery AID-mediated UPF1 depletion RNA-Seq datasets. Transcripts were selected as upregulated when at least one timepoint was significant compared to control 0h IAA condition (padj < 0.0001 & log2FC > 1). Number of transcripts per cluster are indicated. (**C**) Barplot showing the proportions of GENCODE-annotated transcripts per indicated biotype classified as up- or downregulated, complex (= identified in both up- and downregulated clusters), expressed without significant changes (n.s. = not significant) or found as not expressed. (**D**) As in (C) proportions of transcripts per GENCODE biotype identified in individual significantly regulated differential transcript expression (DTE) cluster are shown. (**E**) UpSet plot of the overlap of significantly up- or downregulated transcripts (padj < 0.0001 & abs(log2FC) > 1) between AID- and FKBP-mediated UPF1 depletion for 12h. (**F**) Barplot showing the number of transcripts upregulated in all three UPF1-depletion conditions from (E) per DTE cluster and stratified by GENCODE biotype. (**G**) Schematic overview of tested transcript properties. (**H**) Correlation between transcript properties and expression changes, showing the coefficient of determination (R2) as heatmap. (**I**) Transcript properties as boxplot per DTE cluster. Statistical significance was determined using two-sided Dunn’s test of multiple comparisons with the not significant (n.s.) expressed cluster as reference and the Wilcoxon effect size was calculated. (**J**) (Left) differential transcript usage (DTU) analysis of AID- and FKBP-mediated UPF1 depletion/recovery RNA-Seq datasets, showing the number of significantly regulated mRNA isoforms as well as median delta isoform fraction (dIF) values per GENCODE transcript biotype. (Middle) Analysis of alternative splicing on the intron cluster and individual splice site level. (Right) Changes in isoform usage for SR family genes were summarized per GENCODE transcript biotype (sum(dIF)).

### The majority of UPF1-regulated genes and transcripts are NMD-targets

Having access to an extensive dataset derived from two degron systems and two cell lines, along with additional analyses, provides the opportunity to define a robust set of genes regulated by UPF1, which can be utilized for further analyses. To identify the core set of UPF1-regulated genes, we employed a set of criteria that were designed to capture genes that are commonly regulated in both the AID and FKBP degron systems, as well as genes that are commonly regulated in HCT116 and HEK293 cell lines. We categorized these genes into cloud, shell, or core sets based on their identification as significantly regulated across conditions after 12h UPF1 depletion as follows: core genes are consistently regulated in all three conditions, shell genes are found in two out of three conditions, and cloud genes were identified in only one of these conditions (Table S6). The core of upregulated genes was more than ten times larger (n=1151) than that of downregulated genes (n=92), thus confirming UPF1’s role as an RNA helicase involved in mRNA degradation (Figure 6A). In addition, we next sought to identify genes where changes in RNA expression levels could be attributed to reduced degradation, as indicated by SLAM-Seq data. Notably, the core contained the largest fraction of genes exhibiting stabilization (37%, n=425), followed by the shell and the cloud. Finally, these transcripts were cross-referenced with Ribo-Seq data to confirm the presence of concordant alterations in translation rates. The rigorous selection process resulted in 188 bona fide UPF1-regulated genes that met all criteria, with the majority being coding genes. The list of UPF1-regulated genes is expected to include both NMD-targeted mRNA as well as mRNA controlled by alternative degradation pathways. Therefore, we next aimed to distinguish between these different pathways. To achieve this, we compared the core of UPF1-regulated genes with genes regulated by other NMD factors, utilizing datasets from depletions of SMG5/6/7 and UPF3 (A and B) (Figures S6A, C). Similar to the categorization we made for UPF1, these datasets were used to define core sets, and comparisons were made between the cores of UPF1, SMG5/6/7, and UPF3 (Figure 6B). The overlap of all three cores is expected to primarily encompass NMD substrates, totaling 355 genes (31%), and is therefore labeled as high-confidence NMD targets. The intersections of UPF1 with a single core, either SMG5/6/7 (n=275) or UPF3 (n=118), are denoted as medium-confidence NMD targets (34%) (Figures S6B, D). The remaining genes in the UPF1 core exhibited no overlap with the cores of other NMD factors, but some displayed overlaps with their shells or clouds. Based on the overlap, these were categorized as low-confidence NMD targets or minimal-confidence NMD targets. 97 genes from the UPF1 core that were not found in any intersections with NMD factors are designated as non-NMD targets (Figure 6C). Comparison of the different confidence categories with a published NMD meta-analysis revealed decreasing NMD significance with lower confidence targets (Figure 6C) ^65^. However, more than half of the designated non-NMD targets were still significantly dependent on SMG6 and/or SMG7 (Figure 6C), suggesting that even the non-NMD targets may still contain mRNAs degraded by NMD. This is also reflected by the low, but detectable upregulation of this class of targets in the SMG5/6/7 and UPF3 data. Combining the NMD confidence levels with the previously defined 188 bona fide UPF1-regulated genes allows further classification and identification of e.g. 57 high-NMD-confidence genes that are commonly upregulated, stabilized more translated (Figure 6D). Among those genes are many well-known NMD-targeted genes, such as GADD45B, GAS5, and ZFAS1, indicating that our selection criteria successfully capture bona fide NMD targets. UPF1 was described to play essential roles in several other decay pathways, including Staufen-Mediated mRNA Decay (SMD), which degrades RNAs with Staufen (STAU) binding sites ^41^. To test the role of UPF1 in SMD, we overlapped the UPF1 core, shell, and cloud genes with previously identified RNAs that were found via hiCLIP to interact with STAU1 ^101^. Although the overall overlaps were rather low (in total 380 genes), we detected 94 core-UPF1 genes to be STAU1-binding targets, of which 54 were stabilized by UPF1 depletion (Figure S6E). However, 49 of these genes were also found in SMG5/6/7 and UPF3 data (= minimal to high NMD confidence), suggesting that NMD is responsible for or at least contributes to their degradation. We also cross-correlated the identified hits with recently published STAU1-KO RNA-Seq data from HCT116 cells and found no substantial evidence for upregulation ^102^ (Figure S6F). These results are consistent with a previous study reporting that only a small proportion of UPF1 targets are regulated by STAU1 ^103^.

**Figure 6.**
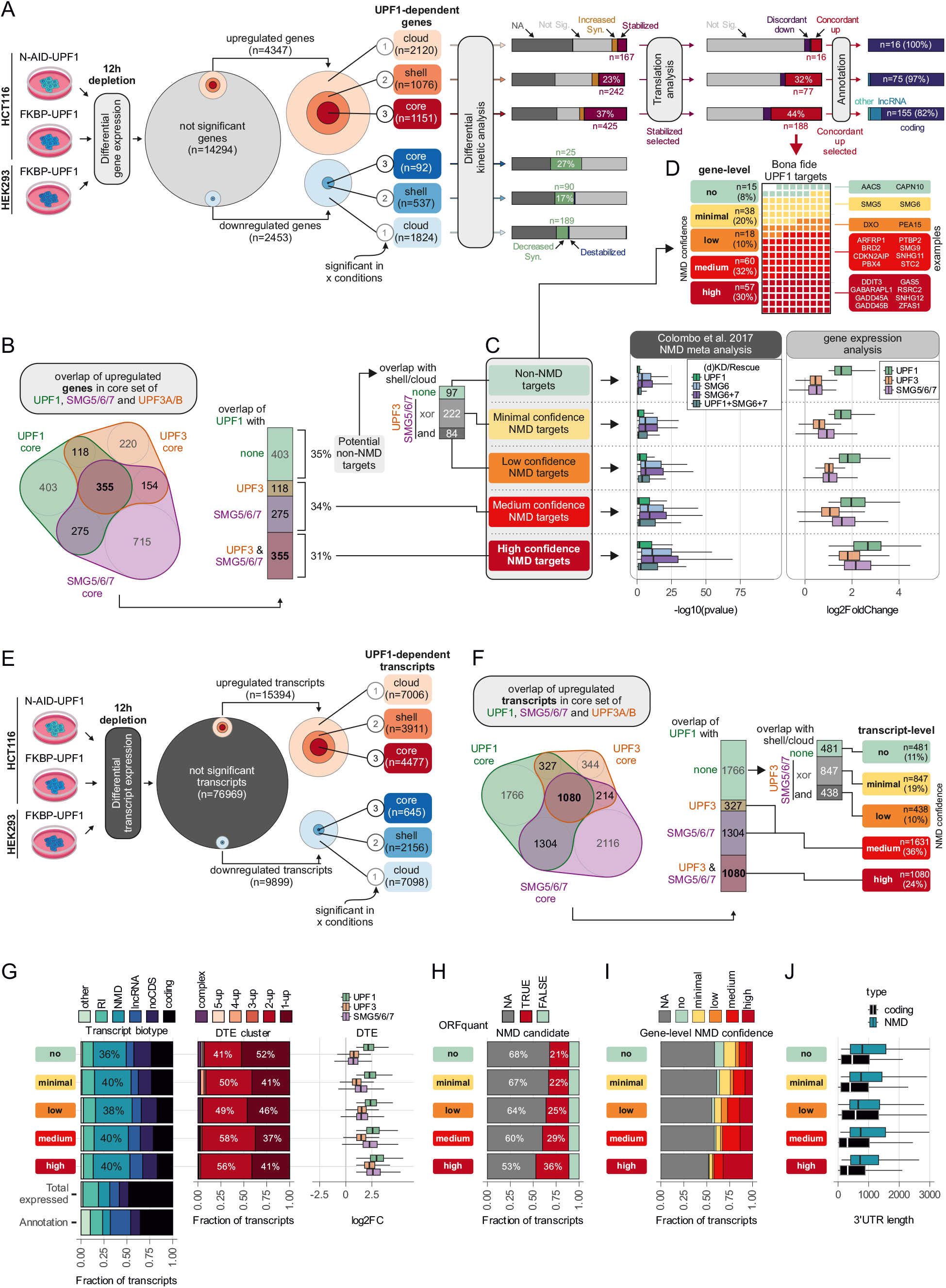
Definition of core UPF1-regulated genes and transcripts. (**A**) Genes with significant expression changes upon 12h UPF1 depletion were categorized in core, shell and cloud based on the number of overlaps between N-AID-UPF1 or FBKP-UPF1 conditions. Further characterization using differential kinetic analysis (SLAM-seq), translation analysis (Ribo-Seq) and classification into GENCODE biotypes was performed. 189 bona bide UPF1 targets were identified. (**B**) Overlaps between core upregulated genes of UPF1, SMG5/6/7 and UPF3A/B conditions. (**C**) Classification of core genes into NMD confidence levels and depicting their statistical significance from published NMD meta analyses (left) or gene expression changes in the indicated datasets (right). (**D**) Combination of bona fide UPF1 target genes and NMD confidence levels, highlighting individual example genes. (**E**) Transcripts with significant expression changes upon 12h UPF1 depletion were categorized in core, shell and cloud based on the number of overlaps between N-AID-UPF1 or FBKP-UPF1 conditions. (**F**) Overlaps between core upregulated transcripts of UPF1, SMG5/6/7 and UPF3A/B conditions. (**G**) Proportions of transcripts per NMD confidence level and GENCODE transcript biotype (left), as well as their categorization into DTE cluster (middle) and expression changes (right) are shown. (**H**) Proportions of predicted ORFs per NMD confidence level and predicted NMD status by applying the 50-nt rule. (**I**) Proportion of transcripts matching the corresponding gene-level NMD confidence. (**J**) 3′ UTR length per NMD confidence level and biotype.

Next, we analyzed the involvement of UPF1 in Histone-Mediated mRNA Decay (HMD). As expected, many histone mRNAs were not sufficiently captured in our poly(A)-selected RNA-Seq, as these transcripts lack a poly(A)-tail (Figure S6G). Therefore, we re-sequenced control, 12h, and 48h IAA-treated N-AID-UPF1 samples using Ribo-minus library preparation, allowing the detection of 73 out of 118 HGNC-annotated histone genes (Figure S6G). Of these genes, only three were significantly upregulated (H2BC1, H2AB2 and H3-4) in 12h or 48h UPF1 depletion (Figure S6H). Unexpectedly, most histone genes exhibited lower expression levels, contradicting the anticipated general role of UPF1 in histone degradation. However, these results are compatible with UPF1-depleted cells exhibiting altered cell cycles, potentially resulting in fewer cells entering S-phase. Collectively, we find little evidence on the gene-level for UPF1 playing a major role in other RNA degradation mechanisms besides NMD. To explore the UPF1-dependent transcriptome further, we performed the core, shell, cloud categorization analyses also at the transcript level without SLAM-Seq data (Figure 6E and Table S7). In total, the categories with high and medium NMD confidence encompassed over 2700 transcripts, 1080 and 1631, respectively (Figure 6F). For 481 transcripts no evidence of NMD involvement was found, as determined by the lack of upregulation in conditions of UPF3 or SMG5/6/7 depletion (Figure 6F). We next asked which annotated transcript biotypes are captured by the NMD classification on the isoform level. Compared to the complete annotation or all expressed transcripts, NMD represented the largest fraction of biotypes in all five confidence levels (Figure 6G). Even the non-NMD transcript class was comprised of 36% annotated NMD-targeted mRNAs. Nearly all targets of the five classes were previously identified as cluster 1 (“early”) or 2 (“delayed”) and displayed on average increased expression levels in SMG5/6/7 and UPF3-depleted conditions. We then used ORFquant ^104^ on our Ribo-Seq data to detect translated ORFs at the transcript-level, successfully assigning ORFs for about 30-50% of the transcripts. ORFquant allows the prediction of ORFs as NMD candidates based on the distance of the stop codon to the last exon-exon junction. As expected, more NMD-inducing ORFs were found with increasing NMD confidence levels (Figure 6H). Of note, approximately 10% of transcripts across all confidence levels were assigned to ORFs that are not predicted to elicit EJC-dependent NMD. Next, we determined the connection between NMD confidence levels on the transcript- and gene-level. Generally, the confidence levels correlated well between genes and transcripts. However, approximately half of the upregulated transcript were not connected to core UPF1-dependent DGE events (Figure 6I), suggesting that these transcripts might contribute to isoform switches. Finally, when exploring the relationship between transcript-level NMD confidence and 3′ UTR length, we found no striking differences (Figure 6J), which is consistent with a recent study ^20^, showing that the length of the 3′ UTR did not correlate with NMD sensitivity.

### NMD RNA substrate regulation supports biological function

Having established that UPF1 regulates many genes via NMD, we hypothesized that there are relevant biological reasons why these genes are particularly regulated by NMD. Further exploration into the mechanisms and identifying individual features responsible for NMD activation will shed light on the interplay between post-transcriptional regulation and cellular functions. We chose to analyze the most robust targets by selecting the bona fide UPF1-regulated genes with high and medium NMD confidence (total 117 genes as in Figure 6D). After ORF prediction on the transcript isoform-level (Table S8), we determined the length-adjusted relative ORF usage (Figure S7A) and predicted the NMD status by applying the established 50-nt rule. We used a cutoff of 10% translation events aggregated across all ORFs as threshold above which EJC-dependent NMD occurs, which was the case for 86 out of 117 genes (Figures 7A-B). Investigating the predicted ORF type in relation to the GENCODE annotation, we observed that the 20 lncRNAs with predicted ORFs exhibit exclusively unannotated (novel), short and > 90% EJC-NMD-inducing translation events (Figures 7A-C and S7B). This aligns with the function of these lncRNAs as hosts for small RNAs, where the fully processed RNA itself seems to lack a biological function. Interestingly, protein-coding genes with similarly high EJC-NMD prediction (> 90%) were mostly driven by short upstream ORFs (uORFs; 13/20 genes). Among those 13 genes with uORF-translation coupled to NMD are well characterized examples such as DDIT3 (CHOP) and PPP1R15A (GADD34), which play an important role in the integrated stress response (ISR) ^105–108^. Conversely, genes that were predicted to be EJC-independent (< 10% EJC-NMD ORF usage) included GADD45B (Figure S7C), which was previously shown in reporter assays to induce NMD by its GC-rich 3′ UTR^103^. When we cross-matched our predicted ORFs with a recently published standardized catalog of Ribo-seq ORFs^109^, we found 20 unannotated ORFs that were replicated by at least one previous Ribo-seq experiment. To further cross-validate our prediction of EJC-dependency, we used two RNA-Seq datasets of EJC downregulations (CASC3 KO and EIF4A3 KD, Figure 7A). We have recently reported that the knockout of the peripheral EJC component CASC3 impairs NMD but not nuclear functions of the EJC^110^. As comparison, we have generated RNA-Seq data from knockdown experiments targeting the core EJC factor EIF4A3. The majority of bona fide UPF1-regulated genes (77/117) were significantly upregulated in EIF4A3-KD cells, including several genes that were not predicted as EJC-NMD-dependent by the ORF analysis (21/31) (Figure 7B). When also considering the core of upregulated CASC3 KO genes, only 3 of the 21 genes remain, namely two lncRNAs (EPB41L4A-AS1, ENSG00000291078) with no significantly predicted ORFs and the coding gene STX1A. Visual inspection revealed that the STX1A transcript annotation in GENCODE is incorrect and the observed ORF will trigger EJC-dependent NMD. In conclusion, the ORF analysis supported the important role of the EJC as NMD activator, but also identified potentially novel EJC-independent NMD targets. One interesting class of protein-coding genes with very high predicted EJC-NMD ORF usage was the neuroblastoma break-point family (NBPF), of which 5 members were found with high NMD confidence and mostly uORF translation (NBPF8, NBPF9, NBPF11, NBPF12 and NBPF15) (Figure 7B). This family of genes encodes for proteins with up to hundreds of Olduvai domains (earlier called DUF1220), which represent the largest human-specific increase in copy number of any coding region in the genome ^111^. The copy number of Olduvai protein domains correlates with primate brain size ^112^, as well as with disorders of the autistic spectrum ^113^. The NBPF family of genes was found to be expressed during human corticogenesis ^114^, probably driving neural stem cell proliferation ^115^. Notably, we found two additional human-specific genes in our bona fide NMD list that are expressed in the fetal cortex ^114^, the lncRNAs WASH5P and CROCCP2 (Figure 7B), the latter of which was recently shown to promote expansion of cortical progenitors ^116^. Intrigued by these results we first established that known marker genes of corticogenesis ^114^ were not generally upregulated due to UPF1 depletion (Figure S7D). We then asked whether other human-specific genes involved in brain development are misregulated when UPF1 is absent. Surprisingly, we found many duplicated gene family members upregulated in UPF1-depleted cells, including ARHGAP11B ^117,118^, NOTCH2NLA and NOTHC2NLC^114,119^ and TBC1D3 genes ^120,121^ (Figure 7D). These results imply that these evolutionary recently acquired genes with important roles in human brain development are constitutively expressed even in non-neuronal cells, but normally degraded by UPF1 and NMD. In turn, this finding further emphasizes the potential necessity to regulate NMD activity especially during human neurodevelopment (reviewed in ^44,47,122^).

**Figure 7.**
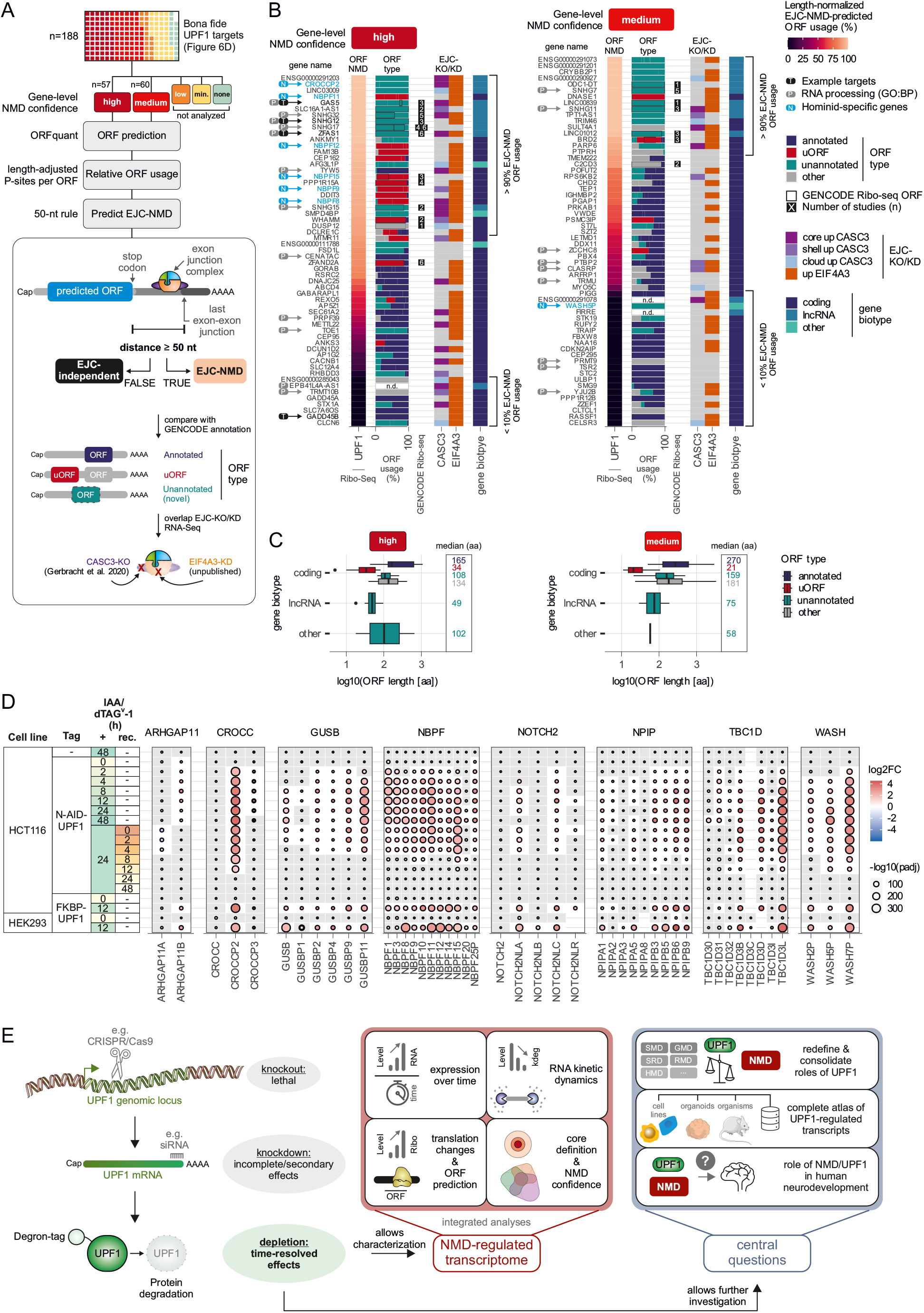
Mechanistic and biological insight into UPF1-regulated genes. (**A**) Workflow overview starting with UPF1 bona fide gene targets, selecting high and medium NMD confidence, using ORFquant to predict ORFs, their relative length-adjusted usage and their EJC-NMD status. ORF type was determined by comparison with the GENCODE annotation and EJC-dependency cross-checked using CASC3 knockout or EIF4A3 knockdown RNA-Seq data. (**B**) All bona fide UPF1 genes with high and medium NMD confidence are shown with their aggregated length-normalized ORF usage (%) that was predicted to elicit EJC-dependent NMD (ORF NMD), in decreasing order. Aggregation was necessary since many genes exhibited multiple predicted ORFs. The proportion of ORF types (annotated, uORF, unannotated [novel] or other) for each predicted ORF is depicted using the length-normalized ORF usage. ORFs found in a previous standardized catalog of GENCODE Ribo-seq ORFs are indicated by the number of studies that identified the same ORF. EJC-dependence is indicated by core/shell/cloud-categorization of CASC3-KO RNA-Seq data or when the gene is upregulated in EIF4A3 knockdown data. Gene biotypes are indicated, as well as target genes previously used as examples, those matching the significant GO:BP term RNA processing and hominid-specific genes. (**C**) ORF length is depicted as boxplot for each ORF type and GENCODE biotype per high or medium NMD confidence. (**D**) Gene expression changes of gene family members implicated in human brain development. (**E**) Model summarizing the key findings of the study. We propose that studying UPF1 protein function is best achieved by using conditional degron tags over classical knockouts (lethal) or RNA interference (incomplete and secondary effects). The strong and time-resolved effects of degron-based approaches allowed the characterization of the NMD-regulated transcriptome by integrating multiple high-throughput analyses. We further envisage that degron-tagged UPF1 allows to investigate key open questions concerning UPF1 and NMD, such as the role in human neurodevelopment.

## Discussion

Among all NMD factors, UPF1 is the most extensively and thoroughly investigated factor by far. PubMed lists more than 650 articles with UPF1 in title or abstract since its initial description in the year 1991^123^ up to the year 2023. While UPF1 is not essential in yeast and C. elegans, providing convenient organisms for knockout studies, the function of UPF1 in higher eukaryotes and mammals has relied on the use of dominant-negative mutants or siRNA-mediated knockdowns ^124,125^. However, these functional genomics approaches, including classical knockouts or RNAi targeting UPF1, suffer from a lack of temporal resolution. In this manuscript, we demonstrate the use of two different conditional degron tags in two different cell lines for the time-resolved analysis of UPF1-regulated gene expression (Figure 7E). The high-throughput data that we obtained provide not only new biological insights but also represent an important resource for the NMD field.

Based on all possible criteria, depleting UPF1 protein via the AID system exhibits functional effects superior to UPF1 knockdowns, even within a short experimental timeframe (8-12 hours). Notably, effects on NMD are observable as early as the initial time point of 2 hours. However, our analysis extended beyond the time points of full NMD inhibition (8/12 hours) and also included longer periods of UPF1 depletion, reaching up to 48 hours. As the duration of UPF1 depletion increased, we observed a greater number and intensity of changes in the transcriptome, including gene and transcript expression changes or the usage of cryptic splice sites. Various analyses indicate that these late occurring changes are primarily due to indirect or secondary effects. Consequently, classical knockouts and prolonged siRNA knockdowns of UPF1 are likely to induce numerous nonspecific gene expression changes, rendering them unsuitable for studies requiring a clear differentiation between specific and nonspecific effects. Another notable technical feature described here is the practically complete reversibility of the rapid UPF1 depletion ^84^. Although the degron cells require a certain amount of time to regenerate sufficient UPF1 protein, shortly after auxin removal the previously upregulated NMD substrates begin to undergo degradation. Our present data do not definitively show whether the identical mRNAs are initially stabilized and subsequently destabilized upon UPF1 repletion. But this assumption aligns with publications suggesting that NMD is possible during each round of translation, even for mRNAs temporarily escaping NMD ^126–128^. In combination with advanced transcriptomics methods, the UPF1 degron cells will provide the means to address this and other questions concerning NMD. The UPF1 degron system, especially the “one-step” dTAG approach, offers versatile applications extending beyond the HEK293 and HCT116 cells utilized in this study. A potential application involves cell types less amenable to siRNA transfection. Additionally, the inducibility of the degrons facilitates inhibition of NMD at specific points in time-dependent experiments.

Our analysis of the UPF1 degron data has enabled us to address important aspects of UPF1’s cellular role. Firstly, we have demonstrated that the majority of mRNAs regulated by UPF1 are genuine NMD targets. This was done by comparing genes/transcripts upregulated by UPF1 with those upregulated by other NMD factors. Most of them are also regulated by at least one other type of NMD inhibition. This analysis leaves some seemingly NMD-independent UPF1-regulated mRNAs that require further investigation. It cannot be ruled out that technical limitations of the used experimental data (sequencing depth, knockdown efficiency, etc.) explain a certain part of these “orphan” UPF1 targets, but they could also be regulated by a UPF1-dependent mechanism other than NMD. This prompted another analysis where we aimed to find evidence supporting previously reported UPF1 pathways in our data. For example, we detected on the transcript-level no global dependence of UPF1 targets on 3′ UTR length, structure, or GC-content. These results question the impact of previously proposed pathways such as structure-mediated RNA decay ^39^, while supporting recent findings debunking long 3′ UTRs as NMD-activating hallmark ^20^. Supporting a previous study ^103^, we also found no indication that STAU1- and UPF1-regulated mRNAs share a substantial intersection. Furthermore, we were unable to identify UPF1-mediated degradation of histone mRNAs. Notably, histone mRNAs showed decreased expression levels following UPF1 depletion, which could be attributed to reduced cell proliferation in the absence of UPF1. However, gaining more detailed insights into these and potentially other UPF1-dependent degradation pathways would require similarly specific and complementary data for the respective pathways, as are available now for UPF1 in our dataset (Figure 7E).

In our final analysis, we explored the biological relevance of the bona fide NMD substrates within our dataset. We observed well-established phenomena, such as NMD’s impact on the integrated stress response and that it targets snoRNA host genes and other lncRNAs (reviewed in ^129^). The two latter classes of RNAs are likely subjected to very efficient cytoplasmic degradation by NMD, given their primary biological role in supplying small RNAs. This idea is supported by our observation that all NMD substrates belonging to this category contain unannotated (novel) ORFs. In this case, NMD acts most likely as a mechanism to eliminate non-functional RNA scaffolds. In other instances, the role of NMD in gene expression is more complex and serves regulatory functions. Among the NMD sub-strates we identify numerous human-specific genes associated with brain development. Further experiments will be required to determine whether NMD facilitates regulation during specific stages of differentiation, as has been postulated in certain contexts ^62,130,131^. Another potential explanation is that the influence of NMD establishes a transcriptional “sandbox”, providing an environment for genes to undergo evolutionary testing that is largely independent of their encoded proteins. The development of the human brain could benefit from this transcriptional plasticity that evolves novel RNA and protein variants. Gaining deeper insights into this fascinating field will require advanced neuronal systems, with the UPF1 degron system being an important molecular tool for these future endeavors.

The bona fide UPF1 targets described in Figures 6 and 7 represent merely a fraction of the NMD-targeted transcripts present in human cells. This is evident from the number of NMD substrates found in the shell and cloud of UPF1-regulated mRNAs. Our rigorous analysis filtered out for example cell-type-specific NMD substrates or those with too few usable reads in the SLAM-Seq analysis. Even among the lower confidence NMD substrates many transcripts exhibit the typical characteristics of NMD-targeted mRNAs either fully or partially. Thus, the dataset is richer in NMD substrates than it may appear at first sight, but we deliberately focused on NMD substrates with the highest confidence during our final analyses. Incorporating additional datasets will further expand the pool of high-confidence NMD transcripts and offer deeper insights into the regulatory mechanisms governing mRNA regulation by NMD. Since the first description of the global effects of UPF1 depletion in human cells, more than 20 years have passed ^124^. Even though a number of interesting insights into the function of UPF1 were achieved back then, the technological differences between these two studies are undeniable. From the examination of two different degrons in two cell lines, it can be inferred that rapid UPF1 depletion surpasses all other means of NMD inhibition. Due to its inherent advantages, it can be excellently combined with other modern molecular biology approaches - such as in our case with SLAM-Seq and Ribo-Seq. Additionally, the dTAG system offers the possibility to deplete UPF1 or inhibit NMD in various systems, from cell culture systems to whole organisms. This will provide insights into the roles of NMD in adult organisms and the application of NMD inhibition in the treatment of genetic diseases.

## Acknowledgements

We thank Juliane Hancke and Silke Modersohn for technical assistance, as well as members of the Gehring and Landthaler labs for discussions and reading of the manuscript. We would like to thank the MDC/BIH Genomics Technology Platform for sequencing. This work was supported by grants from the Deutsche Forschungsgemeinschaft (LA 2941/6-1) to M.L. and (GE 2014/4-1; GE 2014/6-2; GE 2014/13-1) to N.H.G and the Center for Molecular Medicine Cologne (CMMC) [C 05 to N.H.G.].

## Author Contributions

Conceptualization, M.L., N.H.G. and V.B.; Methodology, D.W., N.B., M.L. and N.H.G.; Software, V.B.; Formal analysis, V.B.; Investigation, D.W. (low-throughput assays, sample preparation for RNA-Seq), P.O.W. (Initial Test-RNA-Seq, SLAM-Seq, proteomics), L.G.T.A. (ribosome profiling), M.R. (ribosome profiling), M.F. (RNA sequencing), O.P. (proteomics), J.V.G. (EIF4A3 knock-downs), S.D.G. (SLAM-Seq), K.P. (low-throughput assays) and N.B. (AID-tagged cells); Resources and Data Curation, V.B., K.B., P.M. and E.W.; Writing – Original Draft, Review & Editing, V.B., N.H.G. and M.L.; Visualization, V.B.; Supervision and Funding Acquisition, N.H.G. and M.L.;

## Methods

### RESOURCE AVAILABILITY

#### Lead contact

Further information and requests for reagents should be directed to and will be fulfilled by the Lead Contact, Niels H. Gehring.

#### Materials availability

All cell lines and plasmids generated in this study are listed in Table S1 and are available upon request to the lead contact.

#### Data and code availability

- This study analyzes publicly available data, which are listed below and in the Table S1.
- RNA-Seq data generated in this study have been deposited at BioStudies/ArrayExpress and are publicly available as of the date of publication. Accession numbers are listed in Table S1.
- All original code has been deposited at GitHub (https://github.com/boehmv/2024-UPF1-degron) and is publicly available as of the date of publication.
- Any additional information required to reanalyze the data reported in this study is available from the lead contact upon request.

## EXPERIMENTAL MODEL AND SUBJECT DETAILS

### Cell lines

Flp-In T-REx-293 (human, female, embryonic kidney, epithelial; Thermo Fisher Scientific, cat. no. R78007; RRID:CVCL_U427) cells were cultured in high glucose DMEM with GlutaMAX supplement (Gibco) and HCT116 (human, male, colorectal carcinoma, epithelial; ATCC, cat. no. CCL-247; RRID:CVCL_0291) cells were cultured in McCoy’s 5A (Modified) Medium with GlutaMAX supplement (Gibco) or 2 mM L-glutamine (Gibco). Media were supplemented with 9-10% fetal bovine serum (Gibco) and 1x Penicillin Streptomycin (Gibco). All cells were cultivated at 37°C and 5% CO2 in a humidified incubator. The generation of degron-tagged cell lines is described below and all cell lines are summarized in Table S1.

## METHOD DETAILS

### Protein structure modelling

The AlphaFold ^132^ structure prediction of UPF1 (AF-Q92900-F1-model_v4) was downloaded from the AlphaFold Protein Structure Database (AlphaFold DB, https://alphafold.ebi.ac.uk) ^133^ and overlayed in ChimeraX 1.6.1^134^ with a crystal structure of a UPF1-RNA-ADP:AlF4-complex (PDB DOI: https://doi.org/10.2210/pdb2XZO/pdb) ^135^ using the Matchmaker function to estimate RNA and ADP/ATP binding sites in the AlphaFold model. The final cartoon was generated using CellScape (https://github.com/jordisr/cellscape) ^136^.

### Essentiality and loss-of-function data

The pan-cancer CRISPR Chronos scores (DepMap Public 23Q2+ Score, Chronos) ^52^ were downloaded from https://depmap.org/portal/download/custom/ on 2023-11-16 and the human population data on loss-of-function tolerance from gnomAD v4 (gnomad.v4.0.constraint_metrics.tsv) were downloaded from https://gnomad.broadinstitute.org/downloads on 2023-11-17. Both datasets were analyzed using R version 4.2.2.

### siRNA-mediated knockdowns

Cells were seeded at a density of 2.5 × 10^5^ cells per well in a 6-well plate and reverse transfected with 2.5 µl Lipofectamine RNAiMAX and 60 pmol of the respective siRNAs according to the manufacturer’s protocol. For the combined SMG6+SMG7 knockdown 30 pmol of both SMG6 and SMG7 siRNAs were used. All siRNAs used in this study are listed in Table S1.

### RNA extraction for low-throughput assays

Cells were harvested with 1 ml in-house prepared TRI reagent per 6 well ^137^ and RNA was isolated according to standard protocols. 150 µl 1-Bromo-3-chloropropane (Sigma-Aldrich) was used to induce phase separation and the washed RNA pellet was dissolved in 20 µl RNase-free water by incubating for 10 min on a shaking 65 °C heat block.

### Semi-quantitative and quantitative reverse transcriptase (RT)-PCR

Reverse transcription was performed with 0.75-2 µg of total RNA in a 20 µl reaction volume with 10 µM VNN-(dT)20 primer and the GoScript Reverse Transcriptase (Promega). 40-80 ng cDNA was used as template in end-point PCRs using the MyTaq Red Mix (Bioline) and 0.2 µM final concentration of sense and antisense primer. After 30 PCR cycles, the samples were resolved by electrophoresis on ethidium bromide-stained, 1% agarose TBE gels, visualized by trans-UV illumination using the Gel Doc XR+ (Bio-Rad) and quantified using the Image Lab software (version 6.0.1, Bio-Rad). Probe-based multiplex quantitative RT-PCRs were performed with the PrimeTime Gene Expression Master Mix (IDT), 2% of cDNA per reaction, and the CFX96 Touch Real-Time PCR Detection System (Bio-Rad). PrimeTime qPCR Assays containing primers and probes were purchased from IDT (B2M = Hs.PT.58v.18759587, ZFAS1 = Hs.PT.58.25163607, GAS5 = Hs.PT.58.24767969) and used at 1x final concentration according to the manufacturer’s instruction. Each biological replicate was repeated in technical triplicates and the average Ct (threshold cycle) value was measured. The housekeeping gene B2M (FAM-labelled) Ct values were subtracted from the target (ZFAS1, Cy5-labelled or GAS5, SUN-labelled) values to receive the ΔCt. To calculate the mean log2 fold changes three biologically independent experiments were used. The log2 fold changes are visualized as single data points and mean. All primers used in this study are listed in Table S1.

### Protein abundance and half-live data

The following data from published studies were used to assess protein abundance and half-lives for: HeLa cells (Supplemental Table 134915_1_supp_65961_p3f7xq of publication) ^71^, 29 human tissues (Table EV5 of publication) ^72^ and mouse NIH 3T3 cells (Table S3 of publication) ^73^. All datasets were analyzed using R version 4.2.2.

### Whole proteome analysis

3 × 10^5^ cells were seeded in 24-well plates one day before starting the depletion experiment. UPF1 depletion was induced with 500 µM indole-3-acetic acid (IAA; Sigma-Aldrich; Cat# I5148) at various time points during the experiment (between 0 and 24 hours before harvesting). All cells were harvested at the same time to minimize differences in cell numbers. Cells were washed twice with ice-cold Dulbecco’s phosphate-buffered saline (DPBS; Gibco; Cat # 14190-094) and then scratched in 2 mL of ice-cold DPBS. The cells were collected and pelleted (300 x g; 5 min; 4 °C), washed three times with ice-cold DPBS, transferred to a 96-well plate and frozen at -80 °C until further use.

The samples underwent lysis and digestion in SDC buffer through automated pipetting facilitated by the AssayMAP Bravo robotic system (Agilent Technologies). Briefly, a pellet of about 1 × 10^6^ cells underwent lysis in 2x SDC-buffer comprising 2% (w/v) sodium deoxycholate (Sigma-Aldrich), 20 mM dithiothreitol (Sigma-Aldrich), 80 mM chloroacetamide (Sigma-Aldrich), and 200 mM Tris-HCl (pH 8). Following heating at 95°C for 10 minutes, the lysates underwent enzymatic digestion employing endopeptidase LysC (Wako) and sequence grade trypsin (Promega) at a protein:enzyme ratio of 50:1, with digestion occurring overnight at 37°C.

The resulting peptides were desalted using the AssayMap protocol and subjected to reversed phase liquid chromatography coupled with mass spectrometry (LC-MS). A total of 1 µg of peptides per sample replicate were injected onto an EASY-nLC 1200 system (Thermo Fisher Scientific) for separation applying a 110 min gradient. An Exploris 480 (Thermo Fisher Scientific) instrument was employed for mass spectrometric measurement and operated in data-independent acquisition (DIA) mode. The raw files underwent analysis using DIA-NN version 1.8.1^138^ in library-free mode. An FDR cutoff of 0.01 and relaxed protein inference criteria were applied, and spectra were matched against a human Uniprot database (2022-03), including isoforms, as well as a common-contaminants database. Subsequent downstream analysis was conducted in R version 4.3.2. MaxLFQ normalized intensities were filtered to retain at 3 valid values per group for each protein. For significance determination, the limma package ^139^ was utilized to compute two-sample moderated t statistics. Nominal P-values were adjusted using the Benjamini-Hochberg method.

### RNA half-live data

Supplementary files from published RNA half-life measurements via DRUID in HEK293 cells ^85^ were downloaded (Gene Expression Omnibus (GEO) accession GSE99517; GSE99517_-HEK.4SU.HL.DRUID.one.csv.gz and GSE99517_-HEK.4SU.HL.DRUID.two.csv.gz), gene names converted with Ensembl BioMart ^140^, and mean half-lifes of both replicates calculated.

### NMD meta-analysis data and STAU1 binding sites

Data from a previous NMD factor meta-analysis was downloaded (Supplemental Table S2) ^65^ and matched via Ensembl gene IDs. Similarly, the data from STAU1 hiCLIP were obtained (Supplementary Table 2) ^101^ and matched via Ensembl gene IDs. Histone mRNA analyses The list of histone genes was obtained from the HUGO Gene Nomenclature Committee at the European Bioinformatics Institute (https://www.genenames.org/data/genegroup/#!/group/864), as previously defined ^141^ and matched using Ensembl gene IDs.

### Generation of AtAFB2-expressing HCT116 cells

As parental cells for auxin-mediated degradation, HCT116 cells were generated to stably express the auxin receptor F-box protein AtAFB2 through integration into the AAVS1 safe harbor locus by Cas9/CRISPR-mediated homologous recombination. 3.0 × 10^5^ cells were seeded in a 24-well plate and transfected at 80–95% confluency using 4 µl Lipofectamine 2000 (Life Technologies) and 0.8 µg plasmids. Cells were co-transfected with pSH-EFIRES-P-AtAFB2-mCherry expressing the auxin receptor F-box protein and pCas9-sgAAVS1 at a ratio of 2:1. pSH-EFIRES-P-AtAFB2-mCherry (Addgene plasmid # 129716 ; http://n2t.net/addgene:129716 ; RRID:Addgene_-129716) and pCas9-sgAAVS1-1 (Addgene plas-mid # 129726 ; http://n2t.net/addgene:129726 ; RRID:Addgene_129726) was a gift from Elina Ikonen. After 2 days, the cells were transferred to a 10-cm dish and kept in medium containing 1 µg/mL puromycin (Invivogen, ant-pr) to select for successful AtAFB2 integration. After another 15 days with continuous medium change ever 3-4 days, individual colonies were picked, and transferred to a 96-well plate. Successful integration was tested by genomic PCR using Phire Direct PCR Kit (Thermo Fisher Scientific). Positive clones expressing AtAFB2-mCherry were subsequently expanded and used for subsequent generation of the UPF1-degron cell lines.

### Immunoblot analysis and antibodies

SDS-polyacrylamide gel electrophoresis and immunoblot analysis were performed using protein samples harvested with RIPA buffer (50 mM Tris/HCl pH 8.0, 0.1% SDS, 150 mM NaCl, 1% IGEPAL, 0.5% deoxycholate). For protein quantification, the Pierce Detergent Compatible Bradford Assay Reagent (Thermo Fisher Scientific) was used. All antibodies (see Table S1) were diluted in 50 mM Tris [pH 7.2], 150 mM NaCl with 0.2% Tween-20 and 5% skim milk powder. Detection was performed with Western Lightning Plus-ECL (PerkinElmer) or ECL Select Western Blotting Detection Reagent (Amersham) and the Vilber Fusion FX6 Edge imaging system (Vilber Lourmat).

### Construction of vectors for endogenous tagging

Degron tagging of endogenous loci was conducted by CRISPR–Cas9-mediated HDR. Donor vectors with two homology arms flanking the degron tag, and Cas9 vectors with specific sgRNAs were constructed for N- and C-terminal tagging of UPF1. For the N-terminal donor vector (UPF1 N-term AID in pUC57-Kan), homology arms were constructed by gene synthesis (Biocat) with around 500 bp upstream and downstream of the start codon. Between these homology arms cloning sites (XmnI and BglII) were also introduced, which were later used to insert a N-terminal tagging cassette with a selection marker and degron. The N-terminal tagging cassette [Blasticidin(BSD)/Hygromycin(Hygro)-P2A-V5-MiniIAA7(AID)] was synthesized with flanking restriction sites, XmnI and BamHI (see Fig. 2A) in plasmid pUC57-Kan (Biocat). The C-terminal donor vector (UPF1 C-term AID in pUC57-Kan) was similarly synthesized as the N-terminal donor vector except that the homology arms were flanking the stop codon, which was completely replaced by cloning sites, NheI and SspI. The C-terminal tagging cassette (AID7-V5-P2A-BSD/Hygro) was also synthesized with these flanking restriction sites in the plasmid pUC57-Kan (see Fig. 2A), respectively. To create the final targeting vector (N- and C-terminal), the donor vectors and the tagging cassette vectors were digested at their introduced restriction sites (see above). The restricted N- and C-terminal tagging cassette (BSD/Hygro-P2A-V5-AID and AID-V5-P2A-BSD/Hyg) was ligated to the digested donor vector (UPF1 N-term AID in pUC57-Kan and UPF1 C-term AID in pUC57-Kan), so that the tagging cassette was between the homology arms. This cloning procedure resulted in 4 targeting vectors (UPF1-N-term-AID-V5-P2A-BSD, UPF1-N-term-AID-V5-P2A-Hygro, UPF1-C-term-AID-V5-P2A-BSD, and UPF1-C-term-AID-V5-P2A-Hygro) which were used to tag endogenous loci of UPF1. SgRNAs were synthesized as two unphosphorylated primers, annealed and inserted into BbsI-cut px458-Cas9 vector. pSpCas9(BB)-2A-GFP (PX458) was a gift from Feng Zhang (Addgene plasmid # 48138 ; http://n2t.net/addgene:48138 ; RRID:Addgene_48138). Information about endogenous targets, HDR templates, primers for HDR templates and sgRNAs is provided in Table S1.

### Generation of homozygously tagged cell lines

HCT116 cell-lines expressing AtAFB2 were used to create the UPF1-degron cell lines. The transfection was similar to the generation of HCT116 + AtAFB2 cell-line (see above). A cloned px458-Cas9 + sgRNA plasmid was co-transfected with two homology arm plasmids containing the mAID7-BSD/Hygro-resistance. Two days after transfection, the cells were transferred to a 10-cm dish containing 100 µg/mL hygromycin B Gold (Invivogen) and 5 µg/mL blasticidin (Invivogen) to select for successful AID7 (degron) integration. After two to three weeks of selection colonies were picked and transferred to 96 well plate. For successful biallelic integration clones were tested by genomic PCR using primers binding in the BSD/Hygro resistance sequence and the genomic region upstream of the homology arm (see “Oligonucleotides”). Two single clones were selected for each target. The functionality of these cell clones was tested by inducing the degradation of the AID-tagged UPF1 by treatment with 500 µM indole-3-acetic acid (IAA; from 500 mM stock in H2O; Sigma-Aldrich, cat. no. I5148) for 24h and subsequent Western Blot analysis with an anti-V5-HRP-antibody (Invitrogen, 46-0708). The sgRNAs, and primers for genomic PCR are listed in Table S1.

### Generation of knock-in cells using CRIS-PITCh v2

For FKBP12F36V-tagging of UPF1, the knock-ins were performed using the CRIS-PITCh v2 system ^142^ consisting of a pX330-BbsI-PITCh encoding the gene specific sgRNA (5′-CCCGTACGCCTCCACGCTCA-3′) and the pCRIS-PITChv2-PurR-FKBP (UPF1 N) donor plasmid (based on Addgene # 91796). pX330-BbsI-PITCh was a gift from Peter Kaiser (Addgene plasmid # 127875 ; http://n2t.net/addgene:127875; RRID:Addgene_-127875) and pCRIS-PITChv2-dTAG-Puro (BRD4) was a gift from James Bradner & Behnam Nabet (Addgene plasmid # 91796 ; http://n2t.net/addgene:91796; RRID:Addgene_91796). The latter contains two 40 bp-long N-terminal UPF1 microhomologies flanking a puromycin resistance gene, a T2A signal, the FKBP12F36V tag, and a linker region. One day prior transfection 2.5 × 105 Flp-In T-REx-293 or HCT116 were seeded in 6-wells. Per well 1.0 µg of pX330-BbsI-PITCh and 0.5 µg pCRIS-PITChv2 donor plasmid were transfected using a calcium phosphate-based system with BES-buffered saline (BBS). Additionally, 0.1 µg of pCI-maxGFP was transfected as visual control for transfection efficiency. For BBS-transfection, HCT116 cells were temporarily cultured in high glucose DMEM with GlutaMAX supplement (Gibco). Two days after transfection, the cells were transferred to a 10-cm dish. After a total of 3-5 days post transfection the cell culture medium was supplemented with 2 µg/ml or 1.5 µg/ml puromycin (InvivoGen) for the Flp-In T-REx-293 and HCT116 cell, respectively, to select for successful knock-ins. Surviving colonies were picked and genomic DNA was extracted using QuickExtract DNA Extraction Solution (Lucigen) according to manufacturer’s instruction. Correct integration was screened via genomic PCR, followed by Sanger sequencing (Eurofins Genomics). The sgRNAs, and primers for genomic PCR are listed in Table S1.

### High-throughput-sequencing

Eleven publicly available RNA-Seq datasets were obtained and analyzed: I) UPF1 knockdown in HEK293 cells ^77^ (Gene Expression Omnibus (GEO) ^143^ accession GSE109143), II) UPF1 knockdown in HeLa cells ^79^ (Gene Expression Omnibus (GEO) accession GSE152435), III) UPF1 knockdown in HEK293 cells ^74^ (Gene Expression Omnibus (GEO) accession GSE176197), IV) UPF1 knockdown in K562 cells ^76^ (Gene Expression Omnibus (GEO) accession GSE204987), V) UPF1 knockdown in Huh7 cells ^78^ (Gene Expression Omnibus (GEO) accession GSE185655), VI) UPF1 knockdown in HCT116 cells ^80^ (Gene Expression Omnibus (GEO) accession GSE179843), VII) UPF1 knockdown in HepG2 cells ^75^ (Gene Expression Omnibus (GEO) accession GSE162199), VIII) SMG5, SMG6, SMG7 knockout/knockdown in HEK293 cells ^19^ (BioStudies ^144,145^ accession E-MTAB-9330), IX) UPF3A/B double knockout in HEK293 cells ^81^ (BioStudies accession E-MTAB-10716), X) CASC3 knockout in HEK293 cells ^110^ (BioStudies accession E-MTAB-8461) and XI) STAU1 knockout in HCT116 cells ^102^ (Gene Expression Omnibus (GEO) accession GSE138441).

Eight different high-throughput sequencing datasets were generated in this study: #1) Initial Test-RNA-Seq of +/-IAA-treated HCT116 control, N-AID-UPF1 and C-AID-UPF1 cells, #2) RNA-Seq of control and 0-48h IAA-treated N-AID-UPF1 in HCT116 cells, #3) RNA-Seq of N-AID-UPF1 recovery for up to 48h after 24h IAA-treatment in HCT116 cells, #4) RNA-Seq of FKBP-tagged UPF1 in Flp-In T-REx-293 and HCT116 cells +/-dTAGV-1 treatment for 12h, #5) RiboMinus RNA-Seq of control and 12h or 48h IAA-treated N-AID-UPF1 in HCT116 cells, #6) SLAM-Seq of control and N-AID-UPF1 +/-IAA for 0, 12 and 24h, #7) Ribo-Seq of N-AID-UPF1 in HCT116 +/-IAA for 12h and #8) RNA-Seq of EIF4A3 siRNA-mediated knockdown in Flp-In T-REx-293 cells.

For dataset #1, total RNA was extracted by lysing cells with TRIzol Reagent (Invitrogen). Libraries were constructed using the NEBNext Ultra II Directional RNA Library Prep Kit (New England Biolabs) and sequenced on a NovaSeq6000 device (Illumina). For datasets #2-6, cells were lysed using 1 ml in-house prepared TRI reagent per 6 well ^137^. The RNA was extracted and purified using the Direct-zol RNA MiniPrep kit including the recommended DNase I treatment (Zymo Research; Cat# R2052) according to manufacturer’s instructions. For datasets #2-3, libraries were prepared using the Illumina Stranded TruSeq RNA sample preparation kit. Library preparation started with 500ng total RNA. After poly-A selection (using poly-T oligo-attached magnetic beads), mRNA was purified and fragmented using divalent cations under elevated temperature. The RNA fragments underwent reverse transcription using random primers. This was followed by second strand cDNA synthesis with DNA Polymerase I and RNase H. After end repair and A-tailing, indexing adapters were ligated. The products were then purified and amplified (15 cycles) to create the final cDNA libraries. After library validation and quantification (Agilent Tape Station), equimolar amounts of library were pooled. The pool was quantified by using the Peqlab KAPA Library Quantification Kit and the Applied Biosystems 7900HT Sequence Detection System. The pool was sequenced on an Illumina NovaSeq6000 sequencing instrument with a PE100 protocol aiming for 75 million clusters per sample. Libraries for dataset #4 were prepared from 500ng total RNA with the Illumina Stranded mRNA Preparation kit. After poly-A selection (using Oligo(dT) magnetic beads), mRNA was purified, fragmented and reverse transcribed with random hexamer primers. Second strand synthesis with dUTPs was followed by A-tailing, adapter ligation and library amplification (12 cycles) to create the final cDNA libraries. After library validation and quantification (Agilent Tape Station), equimolar amounts of library were pooled. The pool was quantified by using the Peqlab KAPA Library Quantification Kit and the Applied Biosystems 7900HT Sequence Detection System. The pool was sequenced on an Illumina NovaSeq6000 sequencing instrument with a PE100 protocol aiming for 75 million clusters per sample. Libraries for dataset #5 were prepared from 500ng total RNA. Enzymatic depletion of ribosomal RNA with the Illumina Ribo-Zero Plus rRNA Depletion Kit was followed by library preparation with the Illumina® Stranded Total RNA sample preparation kit. The depleted RNA was fragmented and reverse transcribed with random hexamer primers, second strand synthesis with dUTPs was followed by A-tailing, adapter ligation and library amplification (12 cycles). Next library validation and quantification (Agilent Tape Station) was performed, followed by pooling of equimolar amounts of library. The pool itself was then quantified using the Peqlab KAPA Library Quantification Kit and the Applied Biosystems 7900HT Sequence Detection System and sequenced on an Illumina NovaSeq6000 sequencing instrument with an PE100 protocol aiming for 100 million clusters per sample.

For dataset #6, thiol modification was performed as previously described ^86^. Shortly, we mixed 1 µg of RNA with 10 mM Iodoacetamide (Sigma-Aldrich; Cat#I6125), 50 mM NaPO4 (pH 8) and 50% DMSO. The reaction was performed at 50°C for 15 minutes, protected from light. Alkylation was quenched with 20mM DTT. RNA was precipitated by adding 2.5 volumes of 100% EtOH and 1 µg GlycoBlue (Ambion; Cat# AM9515) and incubating -80°C for 30 minutes. After cold centrifugation at 16,000 x g for 30 minutes, the pellet was washed with 75% EtOH, re-centrifuged for 10 minutes, air-dried for 5-10 minutes, and resuspended in 5-10 µl of H2O. Poly(A) mRNA isolation and library preparation was carried out with 500 ng RNA as input using the NEB-Next Poly (A) mRNA magnetic isolation module (NEB; Cat#E7490L) and NEBNext Ultra II Directional RNA library preparation kit (NEB; Cat#E7760L) according to the manufacturer’s instructions.

Generation of dataset #7 is described below in more detail. For dataset #8, cells were harvested with 350 µl lysis buffer from the NucleoSpin RNA Plus kit (Macherey & Nagel; Cat# 740984.50) and RNA was purified according to the manufacturer’s instructions. Using the Illumina TrueSeq Stranded Total RNA kit library preparation was accomplished. This included removal of ribosomal RNA via biotinylated target-specific oligos combined with Ribo-Zero gold rRNA removal beads from 1 µg total RNA input. After depletion of cytoplasmic and mitochondrial rRNA, the remaining RNA was purified and fragmented using divalent cations under elevated temperature. The RNA fragments underwent reverse transcription using random primers. This was followed by second strand cDNA synthesis with DNA Polymerase I and RNase H. After end repair and A-tailing, indexing adapters were ligated. The products were then purified and amplified to create the final cDNA libraries. Next library validation and quantification (Agilent tape station) were performed, followed by pooling of equimolar amounts of library. The pool itself was then quantified using the Peqlab KAPA Library Quantification Kit and the Applied Biosystems 7900HT Sequence Detection System and sequenced on an Illumina HiSeq4000 sequencing instrument with an PE75 protocol.

For datasets #2-5 the ERCC RNA Spike-In Mix (Invitrogen, Cat# 4456740) and for dataset #8 the SIRV Set1 Spike-In Control Mix (Lexogen, Cat# 025.03) were added to the samples before library preparation to provide a set of external RNA controls that enable performance assessment. The Spike-Ins were used for quality control purposes, but not used for the final analyses.

### Computational analyses of RNA-Seq data

For standard RNA-Seq analyses, reads were aligned against the human genome (GRCh38, GENCODE release 42 transcript annotations ^13^ supplemented with SIRVomeERCCome annotations from Lexogen; obtained from https://www.lexogen.com/sirvs/download/) using the STAR read aligner (version 2.7.10b) ^146^. Transcript abundance estimates were computed with Salmon (version 1.9.0) ^147^ in mapping-based mode using a decoy-aware transcriptome (GENCODE release 42) with the options –numGibbsSamples 30 –useVBOpt –gcBias –seqBias. After the import of transcript abundances in R (version 4.3.0) ^148^ using tximport (version 1.28.0) ^149^, differential gene expression (DGE) analysis was performed with the DESeq2 R package (version 1.40.1) ^150^. Genes with less than 10 counts in half the analyzed samples were prefiltered and discarded. The DESeq2 log2FoldChange estimates were shrunk using the apeglm R package (version 1.22.1) ^151^. Differential transcript expression (DTE) analysis was performed using the Swish method from the fishpond R package (version 2.6.2) ^152^ based on 30 inferential replicate datasets drawn by Salmon using Gibbs sampling and imported via tximeta (version 1.18.1) ^153^. Transcripts were pre-filtered using 10 counts per transcript in at least one condition as cut-off. General significance cut-offs, as long as not indicated otherwise, were log2FoldChange > 1 & padj < 0.0001 for DESeq2 DGE and log2FC > 1 & qvalue < 0.0001 for Swish DTE. To assess technical metrics for each RNA-Seq dataset, MultiQC (version 1.14) ^154^ was used to aggregate the output of Salmon (total reads, alignment rate, library type) and Bowtie2 (version 2.5.0) ^155^ (rRNA alignment rate). Non-poly(A) RNA genes were chosen based on published results ^156^ and Swish was used to obtain log10-scaled expression values. Differential splicing was detected with LeafCutter (version 0.2.9) ^96^ with the significance threshold adjusted p-value (p.adjust) < 0.0001. Differential transcript usage was computed with IsoformSwitchAnalyzeR (version 1.18.0) ^157^ and the DEXSeq method ^139,158^. Significance thresholds were |dIF| > 0.1 and adjusted p-value (isoform_switch_q_value) < 0.0001. Clustering of Z-scaled log2-transformed counts (+1 pseudocount) was performed with k-means clustering with k = 5, as determined by elbow method.

## Analysis of SLAM-seq data

Reads were mapped with STAR as described above and used as input for bam2bakR (version 2.0.0) ^87,88^ to detect T-to-C mutations in mapped reads. The counts binomial data frame output was used for further analysis with bakR (version 1.0.0). The ‘Stan’ model was used to estimate mutation rates, followed by using the ‘Hybrid’ implementation to fit the data and perform differential kinetic analysis. The mechanistic dissection of gene expression regulation was performed with bakR combined with DESeq2 using the ‘DissectMechanism’ function. Three layers of basal conclusions were determined: a) degradation (kdeg) rates (significance cutoffs bakR_padj < 0.01 & |L2FC_kdeg| > 1), b) RNA expression levels (significance cutoffs DE_padj < 0.01 & |L2FC_RNA| > 1) and mechanistic conclusion (significance cutoffs mech_padj < 0.01). For final classification the following parameters were used: Stabilized (bakR_padj < 0.01 & L2FC_kdeg < -1 & DE_padj < 0.01 & L2FC_RNA > 1 & mech_padj < 0.01 & Mech_score < 0); Destabilized (bakR_padj < 0.01 & L2FC_kdeg > 1 & DE_padj < 0.01 & L2FC_RNA < -1 & mech_padj < 0.01 & Mech_score < 0); Increased Synthesis (not significant kdeg changes & DE_padj < 0.01 & L2FC_RNA > 1 & mech_padj < 0.01 & Mech_-score > 0) and Decreased Synthesis (not significant kdeg changes & DE_padj < 0.01 & L2FC_RNA < -1 & mech_padj < 0.01 & Mech_score > 0).

### Ribosome profiling

Ribo-Seq libraries were generated as previously described ^159^ with some modifications. N-AID-UPF1 HCT116 cells were treated with 500 µM IAA for 12 hours or were untreated (control). Afterwards, the cells were washed with PBS containing 100 µg/ml cycloheximide (Biochemika) before cell lysis. Samples were lysed in polysome buffer (100 mM Tris-HCl pH 7.4, 750 mM NaCl, 25 mM MgCl2, 1 mM dithiothreitol (DTT), 1% Triton X-100, 100 µg/ml cycloheximide and 25 U/ml Turbo DNase (Thermo Fisher Scientific)) and directly frozen in liquid nitrogen. RNA was then digested with 2400 U/ml RNase I (Ambion) for 45 min in slow agitation at RT and digestion stopped with 640 U/ml SUPERase•In (Thermo Fisher Scientific). Monosomes were purified by size exclusion with Illustra MicroSpin S-400 HR columns (GE Healthcare) and extracted with 3 volumes of Trizol LS (Thermo Fisher Scientific), chloroform and the RNA Clean & Concentrator kit (Zymo Research). 5000 ng of the isolated monosomes were depleted of rRNA using a RiboPool kit for Ribosome Profiling (siTOOLs Biotech). The RNA was separated on a 17% Urea PAA gel, 27–30 nt RNA fragments were eluted from the gel and 5′ extremity phosphorylated with 10 U T4 polynucleotide kinase (New England Biolabs) for 1 h at 37 °C. For library generation, the NextFlex small RNA sequencing kit (PerkinElmer) was used according to the manufacturer’s instructions. Each Ribo-seq library was prepared in triplicate and sequenced on a NovaSeq6000 device (Illumina). Ribo-Seq reads were demultiplexed, adapter-trimmed, UMIs (random 4 bases at the 5′ and 3′ ends) extracted with UMI-tools (version 1.1.2) ^160^, length-filtered (≥16 & ≤40 nts) and mapped with Bowtie2 (version 2.5.0) ^155^ against human rRNA, tRNA, miRNA and snoRNA. Unaligned reads were mapped with STAR as described above and mapped reads deduplicated with UMI-tools. Gene-level counts were computed with featureCounts from the Rsubread package (version 2.14.2) ^161^ and DESeq2 used for differential ribosome occupancy analysis. Translation efficiency was calculated with DESeq2 using the deltaTE method as described ^91^. Classification was as follows: Concordant-up (padj_Ribo < 0.01 & padj_RNA < 0.0001 & L2FC_RNA*L2FC_Ribo > 0 & L2FC_RNA > 1); Concordant-down (padj_Ribo < 0.01 & padj_RNA < 0.0001 & L2FC_RNA*L2FC_Ribo > 0 & L2FC_RNA < -1); Discordant_up (padj_TE < 0.01 & L2FC_TE > 1); Discordant_down (padj_TE < 0.01 & L2FC_TE < -1). Quality control of Ribo-Seq reads was performed with Ribo-seQC (version 0.99.0) ^89^ and ORFquant (version 1.02.0) ^104^ used on condition-merged files for splice-aware quantification of translation. Previously cataloged GENCODE Ribo-Seq ORFs^109^ were obtained from GENCODE (https://ftp.ebi.ac.uk/pub/databases/gencode/riboseq_orfs/data/).

### Analysis of SLAM-seq data

Gene ontology Functional enrichments analysis of gene lists (for SLAM-seq ordered by p.adjust(ksyn)) was performed using g:profiler via the R package gprofiler2 (version 0.2.2) ^162^, using gene ontology biological process (GO:BP) as data source, a list of all expressed/detected genes as custom background, domain scope set to ‘custom’ and with g:SCS multiple testing correction method applying significance threshold of 0.05. If applicable, highly significant terms (p.adjust < 0.001) were clustered using the binary cut method from the R package simplifyEnrichment (version 1.11.1) ^163^.

### Time course analysis

ImpulseDE2 (version 0.99.10) ^92^ was used for the time course analysis of differential gene expression with default parameters. The option to identify transient expression changes was enabled and parameters for impulse and sigmoid models were extracted.

### RNA properties and structure prediction

Most mRNA isoform properties were extracted from the GENCODE annotation and reference genome using R packages. Structure prediction was performed using RNAfold (version 2.6.4) ^164^.

### Data Presentation

Schematic representations and figures were prepared/assembled using CorelDraw 2017. Quantifications and calculations for other experiments were performed - if not indicated otherwise - with Microsoft Excel (version 2311) or R (version 4.3.0) and all plots were generated using IGV (version 2.14.1) ^165^, GraphPad Prism 5, ggplot2 (version 3.4.2) ^166^, ggsashimi (version 1.1.5) ^167^, nVennR (version 0.2.3) ^168^ or ComplexHeatmap (version 2.18.0) ^169^.

## QUANTIFICATION AND STATISTICAL ANALYSIS

Most performed statistical tests are already implemented in the used bioinformatic tools. For differential gene expression (DGE) analysis, p-values were calculated by DESeq2 using a two-sided Wald test and corrected for multiple testing using the Benjamini-Hochberg method. For differential transcript expression (DTE) analysis, p-values were calculated by Swish using a Mann–Whitney Wilcoxon test on inferential replicate count matrices and corrected for multiple testing using q-value approaches. P-values were calculated for differential transcript usage (DTU) analysis by IsoformSwitchAnalyzeR using a DEXSeq-based test and corrected for multiple testing using the Benjamini-Hochberg method. For alternative splicing, p-values were calculated by LeafCutter using an asymptotic Chi-squared distribution and corrected for multiple testing using the Benjamini-Hochberg method. For differential kinetic analysis, p-values were calculated by bakR using a z-test and corrected for multiple testing using the Benjamini-Hochberg method. Other statistical tests performed independently of bioinformatic tools used the R package rstatix. Used cutoffs are described in the figures, figure legends or method section.

## Supplemental Information

**Figure S1.**
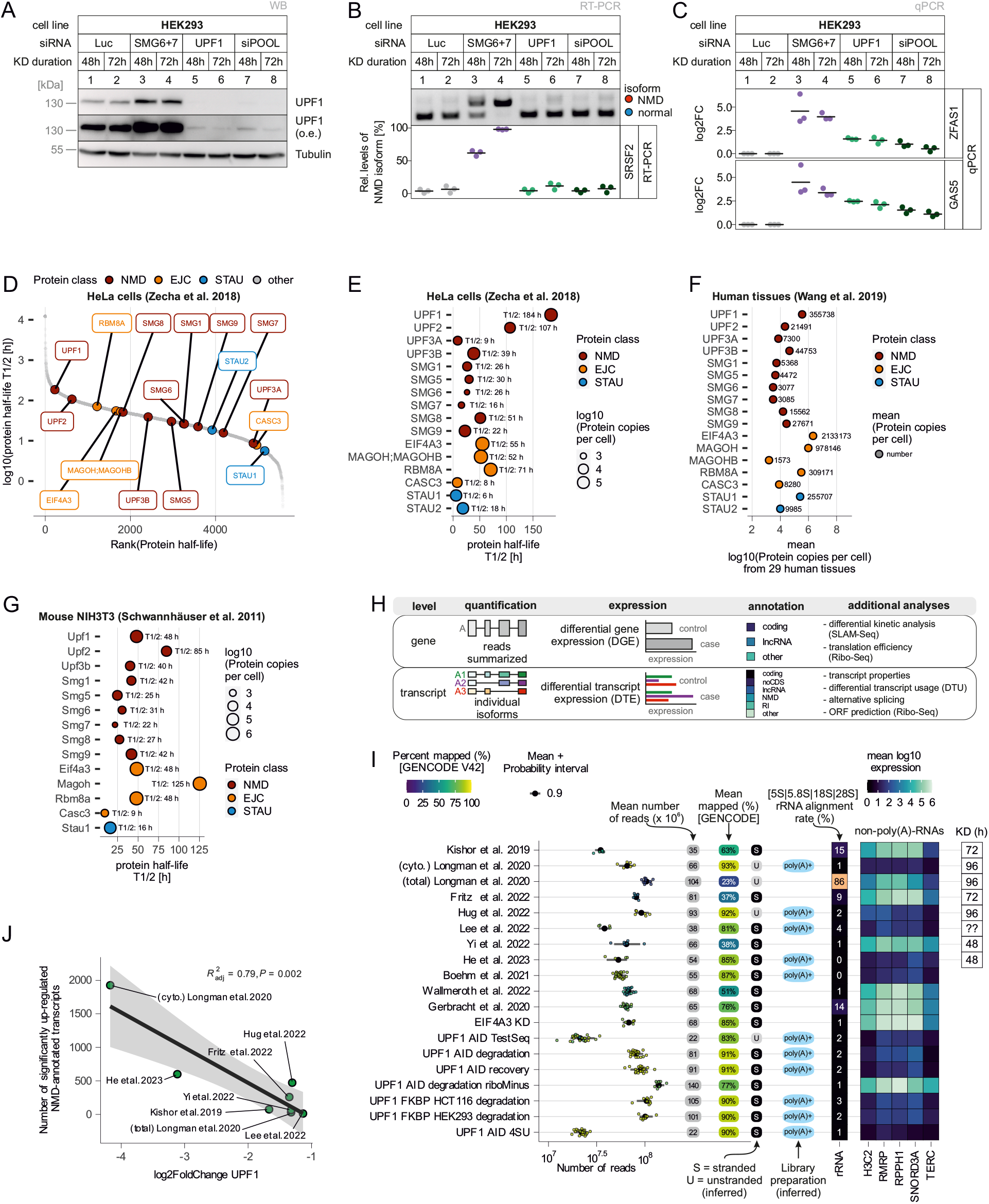
Further characterization of UPF1 downregulation. (**A**) Western blot showing levels of UPF1 protein after knockdown in Flp-In T-REx-293 (HEK293) cells with different siRNAs for 48 or 72h, Tubulin serves as loading control, o.e. = overexposure. (**B**) Detection of the NMD target SRSF2 by end-point RT-PCR in the indicated knockdown conditions in HEK293 cells. The NMD and normal isoforms are indicated and the relative mRNA levels of SRSF2 isoforms were quantified from the agarose gel bands, individual data points and mean are shown (n=3). (**C**) Probe-based qPCR of NMD targets ZFAS1 and GAS5, shown as log2 fold change (log2FC), individual data points and mean are shown (n=3). (**D**) Published data on protein half-lifes from HeLa cells (Zecha et al. 2018) was used to rank proteins based on their log10(half-life) and indicate NMD, EJC and STAU factors. (**E**) Protein half-lives and copy numbers from HeLa cells (Zecha et al. 2018) for NMD, EJC and STAU factors are shown. (**F**) Mean protein copies for NMD, EJC and STAU factors from 29 human tissues (Wang et al. 2019). (**G**) As in (E), but based on data from mouse NIH3T3 cells (Schwannhäuser et al. 2011). (**H**) Overview about gene- and transcript-level high-throughput analyses. For gene-level, RNA-Seq reads are typically summarized from all transcripts isoforms expressed from the particular gene. Differential expression analysis is then performed between control and case conditions based on transcript- or gene-level counts. The GENCODE annotation offers different levels of granularity for transcripts or genes, as for example the NMD-annotation is only available on the transcript-level. Further types of analyses are applicable to genes or transcripts depending on their technical approaches. For example, transcript-level offers precise information about transcripts properties such as length and GC-content. (**I**)Technical characteristics of all RNA-Seq datasets analyzed in this study, including published as well as newly generated datasets. Number of reads per replicate are shown as points, mean reads per dataset are indicated in gray boxes and percentage of mapped reads to GENCODE V42 annotation by Salmon is indicated by fill color. Library strandness was inferred by Salmon and RNA selection method was inferred from mean log10 expression of non-poly(A) RNAs. Also, the percentage of reads mapping to rRNA are shown. For individual published knockdown (KD) datasets, the duration of KD is indicated as declared in the publications. (**J**) Correlation between the gene-level log2FoldChange of UPF1 (determined by DESeq2) and the number of significantly upregulated NMD-annotated transcripts (determined by Swish) for the indicated UPF1 KD RNA-Seq datasets. Linear regression is shown as black line, with confidence interval (0.95) in gray. The adjusted coefficient of determination (adj. R2) and p-value of the correlation is reported.

**Figure S2.**
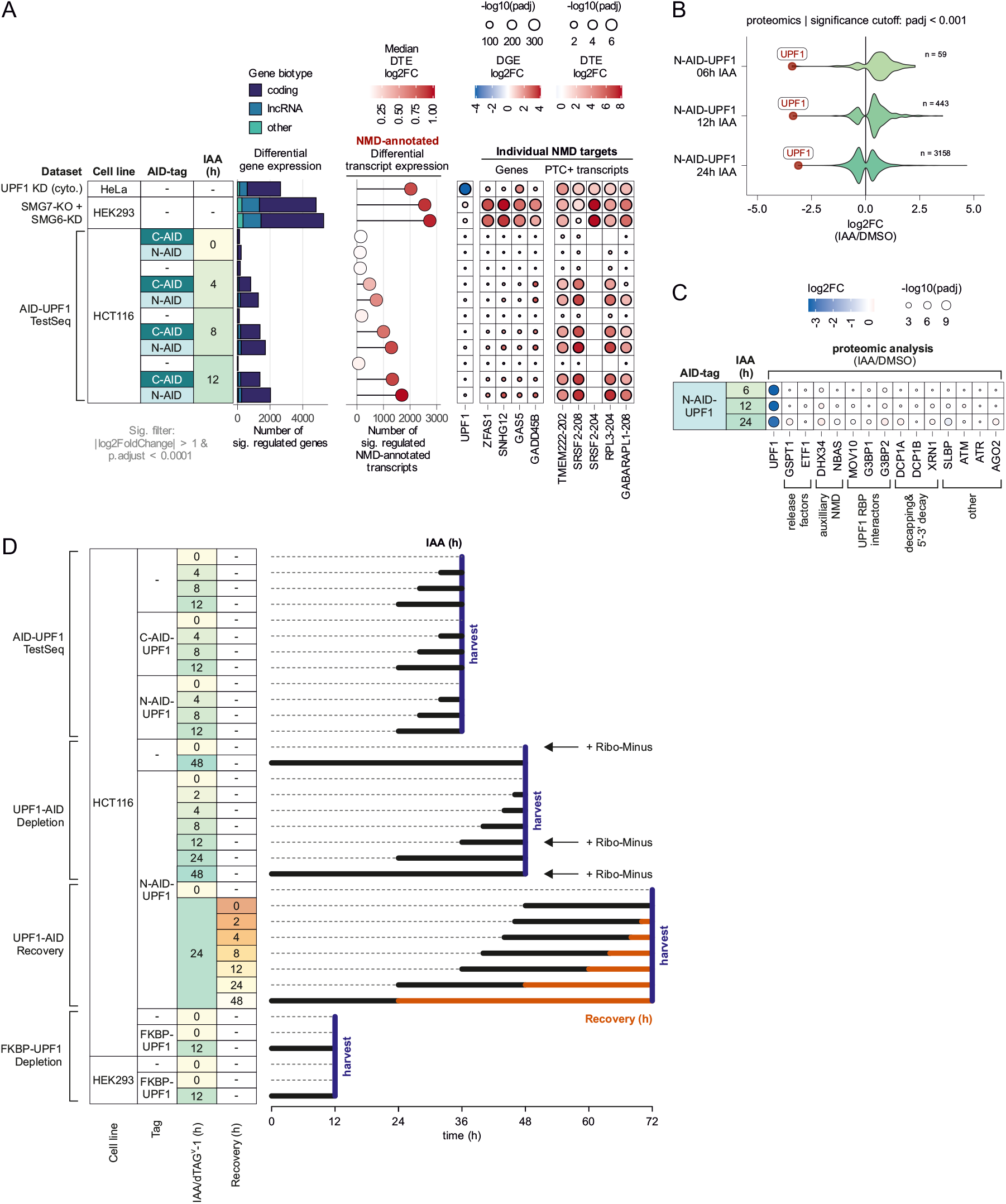
Initial test of AID-tagged UPF1 and experimental overview. (**A**) Comparison of control (parental cells), C-AID-UPF1 and N-AID-UPF1 HCT116 conditions +/-IAA for indicated timepoints with SMG7-KO + SMG6-KD (Boehm et al. 2021) or cytoplasmic UPF1 KD (Longman et al. 2020) RNA-Seq data regarding the number of significantly regulated genes (padj < 0.0001 & abs(log2FC) > 1) stratified by GENCODE biotype (left), the number and median log2FC of significantly regulated GENCODE NMD-annotated transcripts (middle), as well as expression changes of UPF1 mRNA or individual NMD target genes and transcripts (right). (**B**) Violin plot depicting the proteomic analysis of significantly differentially expressed proteins (cutoff padj < 0.001) in the indicated conditions and timepoints, comparing IAA-treated with control cells. The number of significant proteins is indicated as n; UPF1 protein changes are shown as red points. (**C**) Protein expression levels of additional UPF1 interacting factors in IAA-treated versus DMSO control conditions, determined by mass spectrometry. (**D**) Overview of experimental timeline of three datasets, highlighting the timing of IAA addition, if applicable recovery time and the point of sample harvesting.

**Figure S3.**
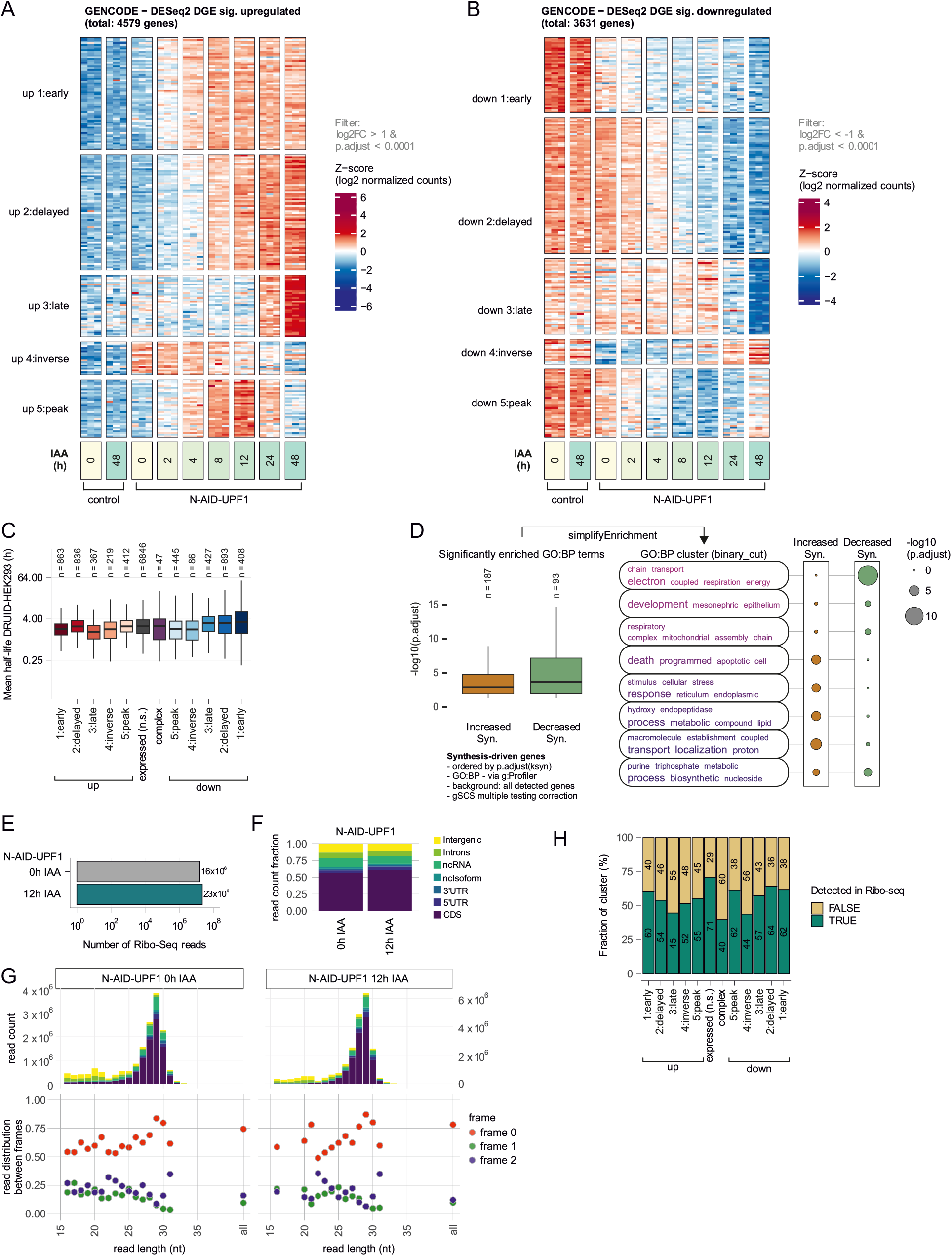
Supporting analyses for the identification of upregulated, stabilized and translated target genes upon UPF1 depletion. (**A**) Heatmap of Z-scores of log2-transformed gene-level counts from upregulated genes (rows) and individual replicates of the indicated conditions (columns), clustered by k-means clustering with k = 5. The total number of genes that were upregulated at least once compared to the 0h IAA control is indicated. (**B**) Heatmap of Z-scores of log2-transformed gene-level counts from downregulated genes (rows) and individual replicates of the indicated conditions (columns), clustered by k-means clustering with k = 5. The total number of genes that were downregulated at least once compared to the 0h IAA control is indicated. (**C**) Boxplot of transcript half-life in hours from DRUID analysis (Lugowski et al. 2018) as mean from two replicates is shown for genes classified into the indicated cluster. Total number of genes per cluster with identified/matchable half-lives is shown. (**D**) Genes detected in the differential kinetic analysis of 12h IAA N-AID-UPF1 compared to 0h IAA control as exhibiting significantly increased or decreased synthesis (Syn.) were ordered by statistical significance of the synthesis rate (p.adjust(ksyn)) and subjected to functional enrichments analysis via g:profiler using the gene ontology biological process (GO:BP). The background gene set were all genes detected in the SLAM-seq analysis. The enriched GO:BP terms are shown with their gSCS-corrected p-value as boxplot, indicating the number of significant terms (n). SimplifyEnrichment was used to cluster the GO:BP terms by binary_cut. (**E**) Number of Ribo-Seq reads aggregated from three replicates for each condition. (**F**) Proportion of Ribo-Seq read counts per condition and aligned biotype or region (e.g. CDS) as determined by Ribo-seQC. (**G**) Read count distribution per read length in nucleotides and biotype/region with the same color scheme as in (F) (top). Frame information per read length (bottom). (**H**) Barplot showing the proportion of genes per DGE cluster that were detected in the Ribo-Seq analysis.

**Figure S4.**
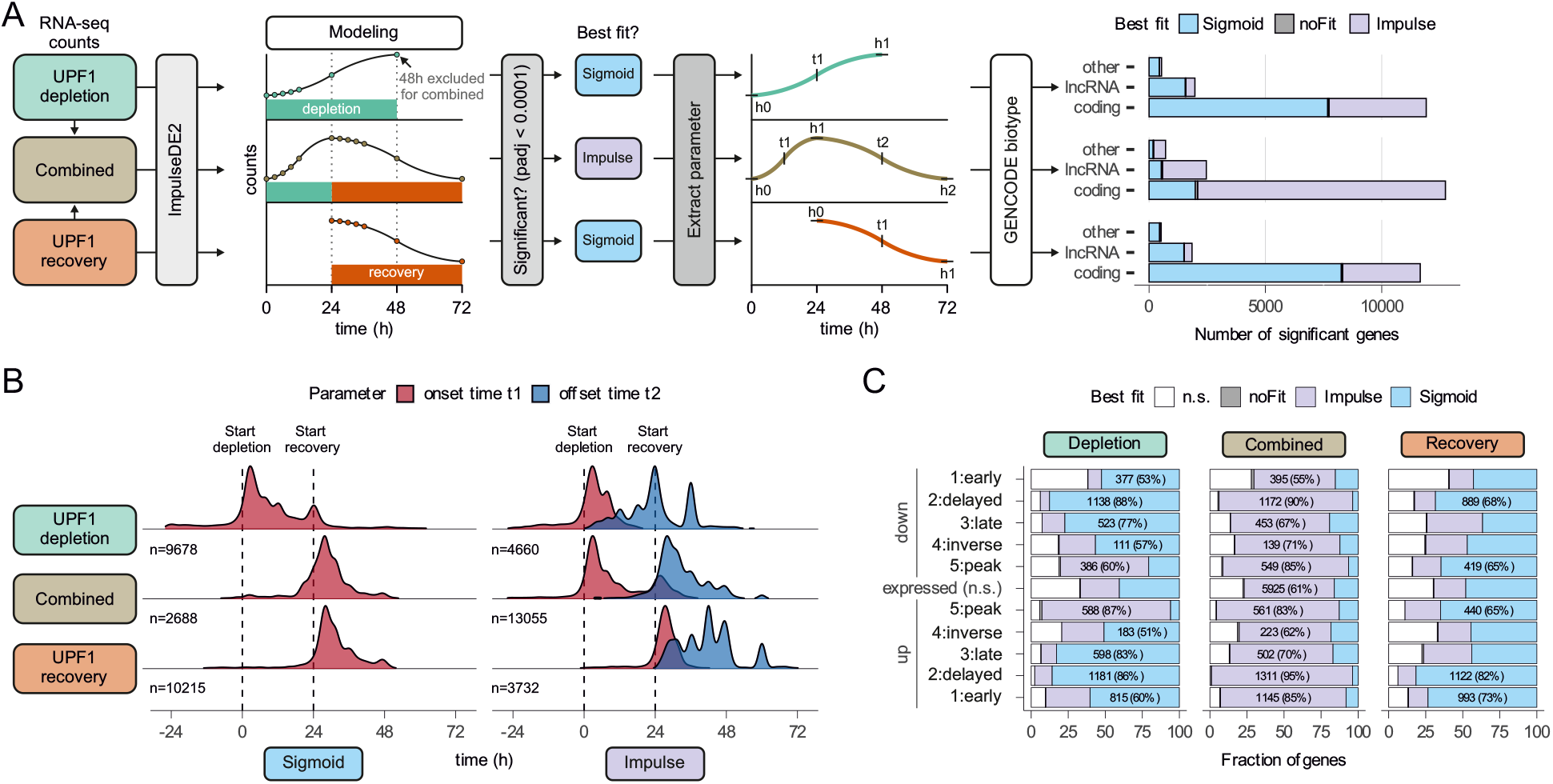
Time course analysis of UPF1 depletion and recovery. (**A**) Schematic overview of the workflow. RNA-Seq counts of UPF1 depletion and recovery were used individually or combined to fit models (sigmoid or impulse model) via ImpulseDE2. For genes with significant time-dependent expression changes the best model was selected and the model parameters were extracted. The number of genes with significant gene expression changes per GENCODE gene biotype and according to their best model fit are shown on the right. (**B**) Density plot showing the distribution of modelled parameters onset time (t1) and offset time (t2) for sigmoid and impulse models per individual or combined dataset. The number of genes is indicated (n). (**C**) Proportion of genes per DGE cluster, best model fit and dataset.

**Figure S5.**
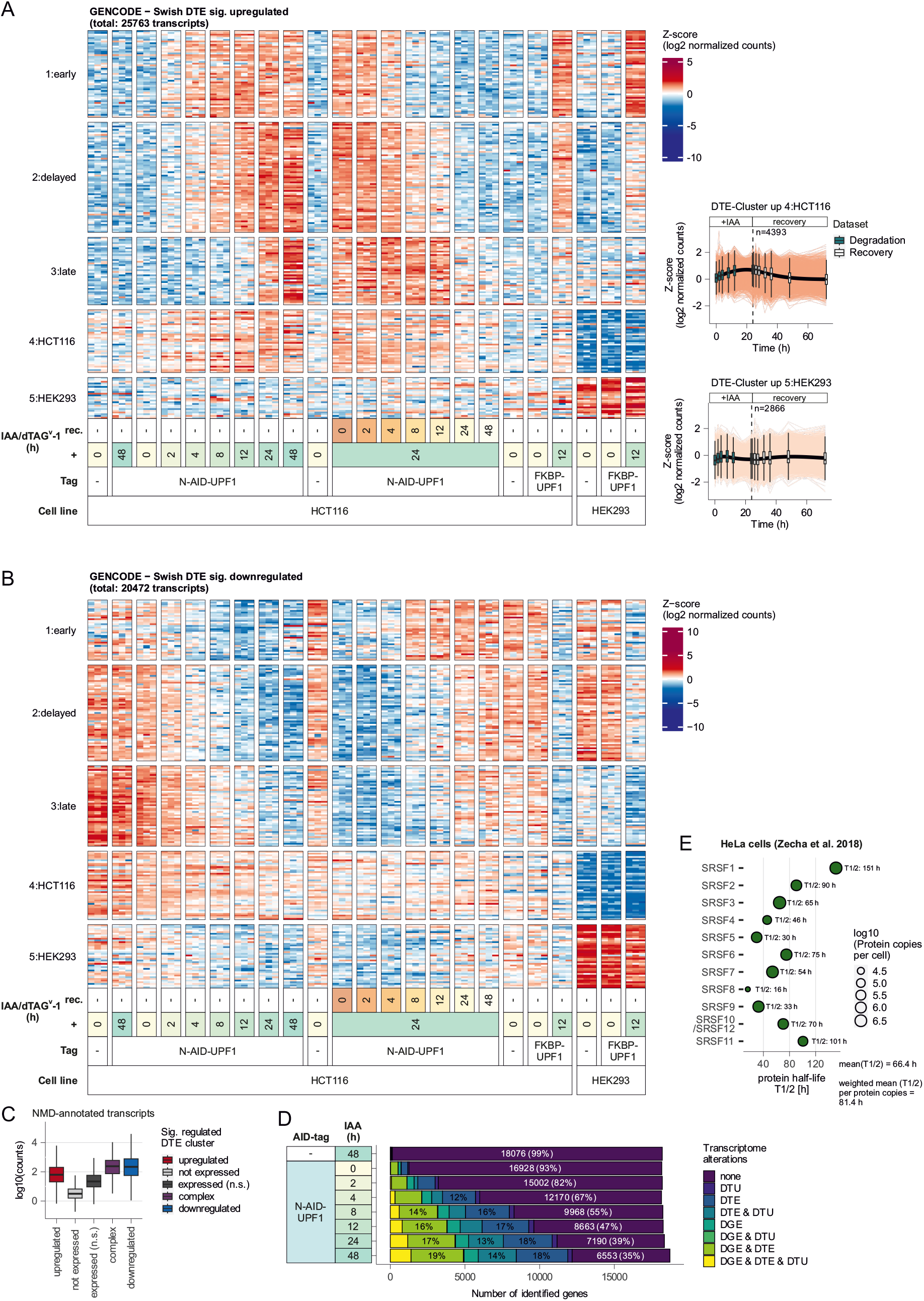
Transcript isoform-specific analyses. (**A**) Heatmap of Z-scores of log2-transformed transcript-level counts from upregulated transcripts (rows) and individual replicates of the indicated conditions (columns), clustered by k-means clustering with k = 5. The total number of transcripts that were upregulated at least once compared to the respective control is indicated. (**B**) Heatmap of Z-scores of log2-transformed transcript-level counts from downregulated transcripts (rows) and individual replicates of the indicated conditions (columns), clustered by k-means clustering with k = 5. The total number of transcripts that were downregulated at least once compared to the respective control is indicated. (**C**) Boxplot of counts of NMD-annotated transcripts per indicated DTE cluster. (**D**) Barplot showing the proportion of genes exhibiting the indicated transcriptome alterations. (**E**) Protein half-lives and copy numbers from HeLa cells (Zecha et al. 2018) for SRSF proteins are shown.

**Figure S6.**
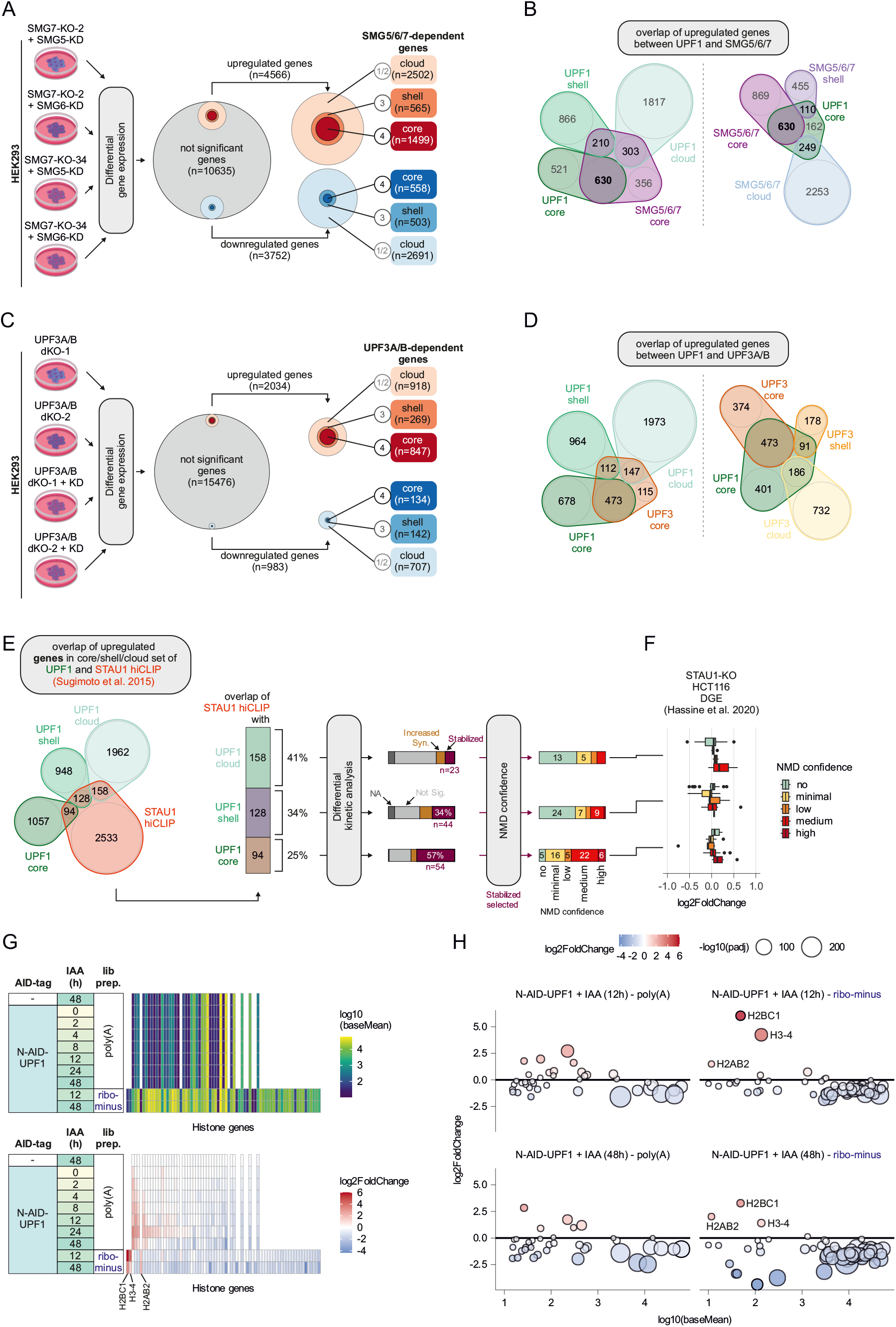
Supporting analyses to UPF1 role in RNA degradation pathways. (**A**) Genes with significant expression changes in SMG7-KO clones (#2 or #34) with additional SMG5 or SMG6 KD were categorized in core, shell and cloud based on the number of overlaps between conditions. (**B**) Overlaps between core upregulated genes of SMG5/6/7 with UPF1 core, shell or cloud sets (left) or between core upregulated UPF1 with SMG5/6/7 core, shell or cloud sets. (**C**) Genes with significant expression changes in UPF3A/B double KO clones (#1 or #2) with or without additional UPF3B KD were categorized in core, shell and cloud based on the number of overlaps between conditions. (**D**) Overlaps between core upregulated genes of UPF3A/B with UPF1 core, shell or cloud sets (left) or between core upregulated UPF1 with UPF3A/B core, shell or cloud sets. (**E**) Overlaps between UPF1 core, shell or cloud sets with STAU1 hiCLIP targets (Sugimoto et al. 2015). Further characterization using differential kinetic analysis (SLAM-seq) and NMD confidence classification. (**F**) Gene expression analysis of identified potential UPF1-STAU1 overlapping stabilized targets in STAU1 KO RNA-Seq data (Hassine et al. 2020). (**G**) Expression levels (top) and gene expression changes (bottom) of histone genes in the respective conditions. (**H**) Combined analysis of expression levels as baseMean from DESeq2 versus expression changes in 12h (top) or 48h (bottom) UPF1 depletion, depending on poly(A) (left) or rRNA depletion (right) library preparation.

**Figure S7.**
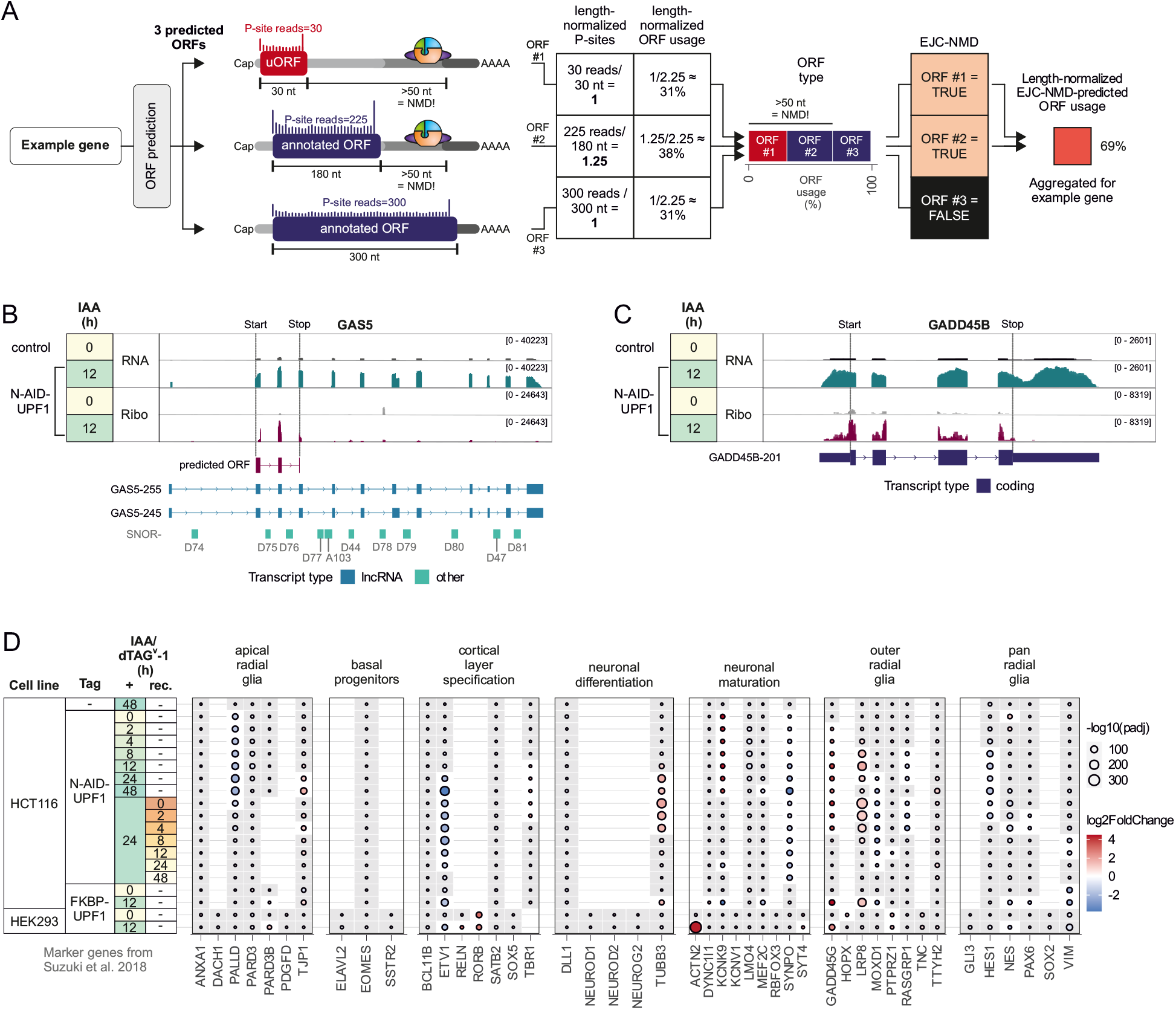
Insights into individual UPF1-targeted transcripts. (**A**) Schematic explanation of the process of determining the length-normalized EJC-NMD-predicted ORF usage of a fictional example gene. Assume ORF prediction with ORFquant yielded 3 ORFs (1x uORF, 2x annotated ORFs) with the indicated number of P-site reads. ORF #1 and #2 are predicted to elicit EJC-dependent NMD since the stop codon is more than 50 nucleotides upstream of the last exon-exon junction. After length normalization of P-site reads, the relative ORF usage is calculated and displayed as stacked bar plot indicating the ORF type. Since ORF #1 and #2 are predicted to induce NMD, their ORF usage is counted for the final analysis. (**B**) IGV snapshot of GAS5 showing read coverage from RNA-Seq and Ribo-Seq from 0h and 12h IAA treatment of N-AID-UPF1 conditions. The predicted ORF is highlighted and the most relevant transcript annotations, including the snoRNAs are depicted below. (**C**) IGV snapshot of GADD45B as in (A). (**D**) Expression changes of genes previously used as marker genes for corticogenesis (Suzuki et al. 2018).

## Supplemental Tables

Table S1. Resource and material lists

Table S2. Differential gene expression analysis, Related to Figure 3

Table S3. Differential kinetic analysis, Related to Figure 3

Table S4. Translation efficiency analysis, Related to Figure 3

Table S5. Time-course analysis by ImpulseDE2, Related to Figure 4

Table S6. Gene-level analyses of UPF1-regulated targets, Related to Figure 6

Table S7. Transcript-level analyses of UPF1-regulated targets, Related to Figure 6

Table S8. ORF prediction by ORFquant, Related to Figure 7

